# PD-1 regulates CD4^+^ T cell-mediated CD8^+^ T cell responses in the brain to balance viral control and neuroinflammation

**DOI:** 10.1101/2025.11.26.690770

**Authors:** Arrienne B. Butic, Elia Afanasiev, Samantha A. Spencer, Mofida Abdelmageed, Anirban Paul, Kalynn M. Alexander, Katelyn N. Ayers, Todd D. Schell, Matthew D. Lauver, Ge Jin, Samantha M. Borys, Laurent Brossay, Jo Anne A. Stratton, Vonn Walter, Aron E. Lukacher

## Abstract

Programmed cell death protein 1 (PD-1) is expressed by T cells during progressive multifocal leukoencephalopathy (PML), a life-threatening brain disease caused by the human-only JC polyomavirus. Why PD-1 blockade finds variable success in PML patients is unclear. Brain CD4^+^ and CD8^+^ T cells are PD-1^high^ during mouse polyomavirus (MuPyV) encephalitis. Here, we show that PD-1 loss during MuPyV infection acts in a brain-autonomous manner to increase the magnitude of brain-infiltrating CD4^+^ and CD8^+^ T cells and the function of virus-specific CD8^+^ T cells; in concert, brain virus levels decline and neuroinflammation increases. Deletion of PD-1 in CD4^+^ T cells, but not CD8^+^ T cells, recapitulates effects of global PD-1 loss. Single-cell RNA sequencing shows that PD-1-deficient CD8^+^ T cells cluster as effectors while transcripts associated with proliferation and function are upregulated with loss of PD-1. Thus, CD4^+^ T cell-intrinsic PD-1 signaling balances antiviral defense against neural injury during polyomavirus CNS infection.

## Introduction

Programmed cell death protein (PD-1) is an immune inhibitory receptor expressed on activated T cells and maintained on precursor exhausted T cells^1^. Binding of PD-1 to either of its two ligands, programmed death-ligand 1 (PD-L1 or CD274) and programmed death-ligand 2 (PD-L2 or CD273), triggers a series of dephosphorylation events that counter costimulatory signals required for T cell activation^1^. With repetitive TCR stimulation, PD-1 signaling directs epigenetic changes that divert T cells toward an exhaustion differentiation state^2^. Disrupting PD-1 signaling enhances the antiviral effector CD8^+^ T cell response to chronic infections in lymphoid tissues and improves the effector activity of tumor-infiltrating CD8^+^ T cells^3,4^. The impact of PD-1 signaling blockade on recovery of exhausted CD8^+^ T cell functionality has been leveraged into the development of immune checkpoint inhibitors (ICIs) that target the PD-1 pathway. Several PD-1 and PD-L1 blocking monoclonal antibodies have dramatically improved survival outcomes for patients with cancers typically characterized by high immunogenicity and/or tumor mutational burden^5^. PD-1 ICIs, however, have minimal benefit for other malignancies and may cause immune-related adverse events (irAEs)^5^.

Although PD-1 inhibition via mechanisms intrinsic to CD8^+^ T cells has been extensively characterized, less is known about inhibition via cell-extrinsic mechanisms. A recent study demonstrated that PD-1-deficient tumor-infiltrating CD8^+^ T cells promoted functionality of PD-1-sufficient CD8^+^ T cells^6^. CD4 ^+^ T cells provide critical help to CD8^+^ T cells during acute viral infection to generate CD8^+^ T cell memory and, during persistent viral infection, maintain CD8^+^ T cell functionality^7,8^. Notably, CD4^+^ T cells express PD-1 during chronic antigen exposure^9^. In the context of cancer, CD4^+^ T cells have been implicated as a systemic requirement for effective CD8^+^ T cell responses after PD-1 blockade^10,11^. Whether PD-1 on CD4^+^ T cells regulates CD8^+^ T cell functionality extrinsic to CD8^+^ T cells is an open possibility.

JC polyomavirus (JCPyV) is a human-specific virus that establishes a lifelong silent infection in the urogenital tract of most healthy individuals^12^. Under immunocompromised conditions, the virus can infiltrate the CNS to cause a variety of life-threatening diseases, the most frequent being the aggressive demyelinating disease, progressive multifocal leukoencephalopathy (PML)^12^. No FDA-approved anti-JCPyV agents exist; current treatment strategies focus on immune reconstitution^13^. CD4^+^ and CD8^+^ T cells in PML patients express high levels of PD-1, raising the potential for PD-1-based immunotherapy^14^. As with cancer immunotherapy, ICI blockade for PML has led to inconsistent outcomes, emphasizing a need to understand how PD-1 influences the anti-polyomavirus immune response in the CNS^15^.

Mouse polyomavirus (MuPyV) is a natural murine pathogen that establishes an asymptomatic persistent infection in immunocompetent mice^16^. Mice inoculated intracerebrally (i.c.) with MuPyV develop a durable population of brain-infiltrating PD-1^high^ CD8^+^ and CD4^+^ T cells^7,17,18^. MuPyV CNS infection in PD-L1-deficient mice induces irAEs in the form of elevated neuroinflammation^17^. Because the CNS is replete with non-renewable neurons, balancing viral control against neuroinflammation-associated pathology is a central concern with PD-1 ICI therapy in the CNS. Using the MuPyV CNS infection model, we asked whether PD-1 intrinsic to antigen-experienced CD4^+^ T cells and/or virus-specific CD8^+^ T cells differentially regulates the antiviral T cell response and neuroinflammation in the brain. We find that PD-1 on CD4^+^ T cells operates extrinsically to limit expansion of brain antiviral CD8^+^ T cells and mitigate virus-induced neuroinflammation at the expense of checking infection.

## Results

### T cells are the major cell type expressing PD-1 in the MuPyV-infected brain

We previously showed that T cells express PD-1 during MuPyV brain infection^17,18^. Although PD-1 functions as an immune checkpoint inhibitor on T cells, it is also expressed by other cell types, such as natural killer (NK) cells^19^ and B cells^20^. To determine whether T cells are the major cell type expressing PD-1 in the brains of MuPyV-infected mice, we utilized multiplexed error-robust fluorescence in situ hybridization (MERFISH) spatial transcriptomics. Wild-type (WT) C57BL/6 mice were i.c. inoculated with MuPyV, and brains were processed for MERFISH at 8 days post infection (dpi), the peak of the anti-MuPyV T cell response^21^. Control mice were sham-injected i.c. with vehicle. Anatomically matched brain sections of cerebral hemispheres from infected mice and sham-injected controls were aligned side-to-side for MERFISH imaging. Cell types were determined as previously described^22^, with ependymal cells defined by double *Cfap6*5 and *Foxj1* positivity as well as high expression of *Foxj1/Cfap6*5 and low expression of *Cd8a/Cd8b1/Csf1r/Cx3cr1*, myeloid cells via double *Csf1r* and *Cx3cr1* positivity, oligodendrocytes through *Mobp* and *Mbp* positivity, oligodendrocyte precursors (OPCs) via double *Pcdh15* and *Pdgfra* positivity, and CD8^+^ T cells through double *Cd8a* and *Cd8b1* positivity. Brain sections of uninfected mice showed no expression of *Pdcd1*, the gene encoding PD-1 (**Supplementary 1A**). In contrast, MuPyV infection recruited a population of CD8^+^ T cells expressing *Pdcd1* **(Supplementary 1B)**. The low-level expression of *Pdcd1* by the ependyma of infected mice is a cell segmentation artefact caused by aggregation of PD-1^+^ CD8^+^ T cells near the ependyma^22^. Because the processed brain sections from infected mice lacked sufficient levels of MERFISH CD4 transcripts for statistical analysis, CD4^+^ T cells were not included in the analysis.

We also utilized MERFISH imaging to ask where PD-L1, the more broadly expressed ligand of PD-1, was expressed. The ependyma lines the cerebral ventricles, constituting a single-cell barrier between the brain parenchyma and the cerebrospinal fluid (CSF), and is the central site of MuPyV replication following i.c. inoculation^22^. In uninfected controls, *Cd274*, which encodes PD-L1, was minimally expressed by ependymal cells of sham-injected mice **(Supplementary 1C)**. During infection, *Cd274* was upregulated by the ependyma **(Supplementary 1D)**. *Cd274* transcripts were also observed in myeloid cells and oligodendrocytes clustered largely around the lateral ventricle as well as in CD8^+^ T cells **(Supplementary 1D)**.

To confirm that PD-1 is predominantly expressed by T cells in the MuPyV-infected brain, WT mice were inoculated i.c. with MuPyV, and lymphocytes were analyzed by flow cytometry at 8 dpi. Virus-specific CD8^+^ T cells were identified with H2-D^b^ tetramers complexed with LT359-368 (LT359), the immunodominant epitope in MuPyV’s Large T Antigen^23^. D^b^LT359 tetramers recognize 50-75% of brain-infiltrating CD8^+^ T cells^17,18,21^. Nearly all antigen-experienced (i.e., CD44^hi^) CD4^+^ T cells in the brains of MuPyV-infected B6 mice recognize one of two viral epitopes^7^. PD-1 was highly expressed by CD44^hi^ CD4^+^ T cells and virus-specific CD8^+^ T cells relative to other cell types examined **(Fig. 1A)**. PD-1 expression on T cells was maintained at similar levels in brains of persistently infected mice (30 dpi) **(Supplementary 2A)**. Circulating T cells in the blood did not express PD-1 at either 8 or 30 dpi, indicating that sustained PD-1 expression on T cells is specific to the brain **(Supplementary 2B)**. Although MERFISH violin plots for *Cd274* indicate that a relatively low proportion of myeloid cells and oligodendrocytes expressed *Cd274* **(Supplementary 1D)**, PD-L1 was highly expressed on CD11b^+^ CD45^high^ macrophages and activated microglia, which upregulate CD45^24^ **(Fig. 1B)**. The few *Cd274*-expressing myeloid cells and oligodendrocytes detected by MERFISH may reflect the high fraction of *Cd274-*negative myeloid cells and oligodendrocytes **(Supplementary 1D)**, tissue sampling bias compared to flow cytometric analysis of PD-L1^+^ cells from whole brain homogenates (**Fig. 1B**), or low RNA copy numbers versus protein levels.

**Figure 1.**
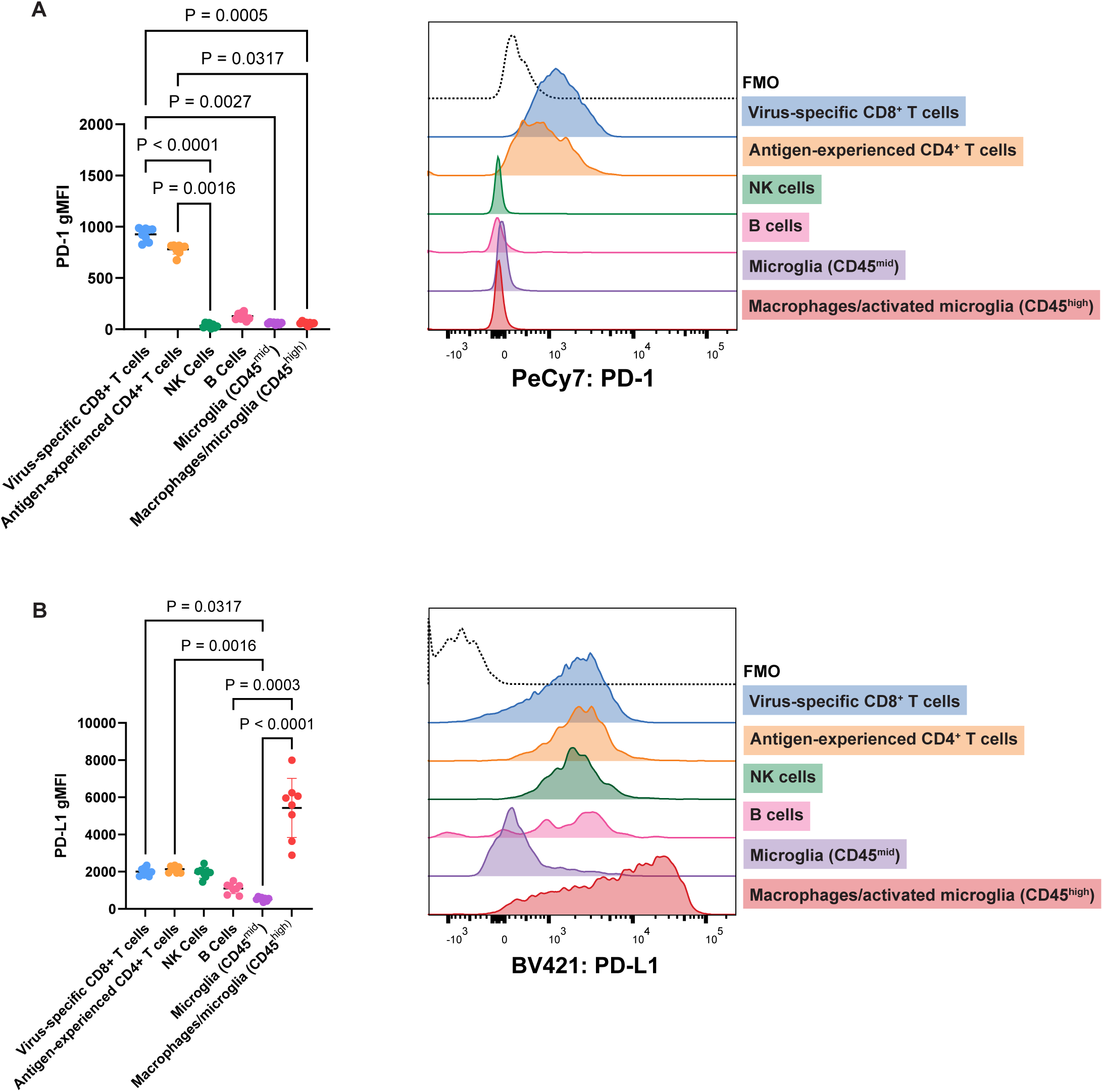
T cells express PD-1, and multiple resident and infiltrating cell types express PD-L1 in the MuPyV-infected brain. WT B6 mice were infected i.c. with MuPyV, and at 8 dpi, cells were isolated from brains and analyzed for PD-1 and PD-L1 expression. Virus-specific CD8^+^ T cells were CD45^+^, CD3^+^, CD8^+^, CD44+, and LT359^+^. Antigen-experienced CD4^+^ T cells were CD45^+^, CD3^+^, CD44^+^, and CD4^+^. NK cells were CD45^+^ CD3^-^ NK1.1^+^. B cells were CD45^+^ CD3^-^ CD19^+^. Microglia were CD3^-^ CD11b^+^ CD45^mid^, and macrophages/microglia were CD3^-^CD11b^+^ CD45^high^. A) PD-1 expression in different cell types in WT brains, shown through geometric mean fluorescence intensity (gMFI) and representative histogram from one mouse. B) PD-L1 expression in different cell types in WT brains, shown similarly as A. n = 8 mice. Error bars show mean ± SD. Data are representative of two independent experiments. Statistical significance for all data calculated with the Friedman’s test, a nonparametric alternative to the repeated measures one-way ANOVA, with a Dunn’s post-test. A repeated measures approach is appropriate because each cell type shown is from one group of mice, rendering the groups dependent on each other.

PD-L1 expression on CD11b^+^ CD45^high^ macrophages and activated microglia is reduced by 30 dpi **(Supplementary 2C)**. In line with and expanding on the MERFISH data **(Supplementary 1D)**, PD-L1 was also expressed by virus-specific CD8^+^ T cells, antigen-experienced CD4^+^ T cells, B cells, and NK cells, albeit at lower levels than on CD11b^+^ CD45^high^ macrophages and activated microglia (**Fig. 1B; Supplementary 1D)**. Together, flow cytometry and spatial transcriptomics demonstrate that during MuPyV infection, PD-1^hi^ CD4^+^ and CD8^+^ T cells cluster around the lateral ventricles in proximity to PD-L1-expressing ependymal cells, activated microglia, and macrophages.

### PD-1 alters the magnitude, function, and differentiation of brain T cells and limits viral control

To investigate whether PD-1 affects T cell function and differentiation during MuPyV encephalitis, we crossed PD-1^fl/fl^ mice with Rosa26-Cre^ERT2^ (Rosa-Cre) mice to generate PD-1^fl/fl^ Rosa-Cre^ERT2^ (PD-1^fl/fl^ Rosa-Cre) mice, in which exon 2 of *Pdcd1* is conditionally deleted after administration of tamoxifen. Cre-negative Rosa-Cre mice were utilized as controls instead of PD-1^fl/fl^ mice to exclude potentially confounding effects of tamoxifen-induced Cre on immune responses.^25^ To verify knockdown of PD-1, PD-1^fl/fl^ Rosa-Cre and control Rosa-Cre mice were injected i.c. with MuPyV, then administered tamoxifen intraperitoneally (i.p.) from 8 to 12 dpi **(Fig. 2A)**. At 15 dpi, PD-1 expression was reduced on both antigen-experienced CD4^+^ T cells and virus-specific CD8^+^ T cells **(Supplementary 3A-B)**. Notably, antigen-experienced CD4^+^ and virus-specific CD8^+^ T cell numbers were both significantly higher in the brains of PD-1^fl/fl^ Rosa-Cre mice than in Rosa-Cre controls **(Fig. 2B)**. No differences in cell counts were exhibited by T cells in the spleens of tamoxifen-treated PD-1^fl/fl^ Rosa-Cre mice **(Fig. 2C)**, consistent with prior work showing that splenic CD8^+^ T cells do not upregulate PD-1 in response to MuPyV infection^18^ and further confirming that changes in T cell differentiation and function with PD-1 loss were relegated to T cells in the brain. T cell counts in the brain exhibited a higher proportion positive for the Ki67 proliferation marker in PD-1^fl/fl^ Rosa-Cre mice **(Fig. 2D)** and no change in the frequency of cells undergoing early apoptosis (Annexin V^+^ Fixable Viability Dye^-^) **(Supplementary 4A-B)**. In addition to acquiring higher proliferative functionality, an increased frequency of virus-specific CD8^+^ T cells was positive for perforin and granzyme B upon *Pdcd1* disruption **(Fig. 2E-F)**. Loss of PD-1 did not lead to an increase in IFN-γ^+^ CD8^+^ T cells, implying that PD-1-deficient CD8^+^ T cells are predominantly deploying a cytotoxic effector mechanism **(Fig. 2G)**.

**Figure 2.**
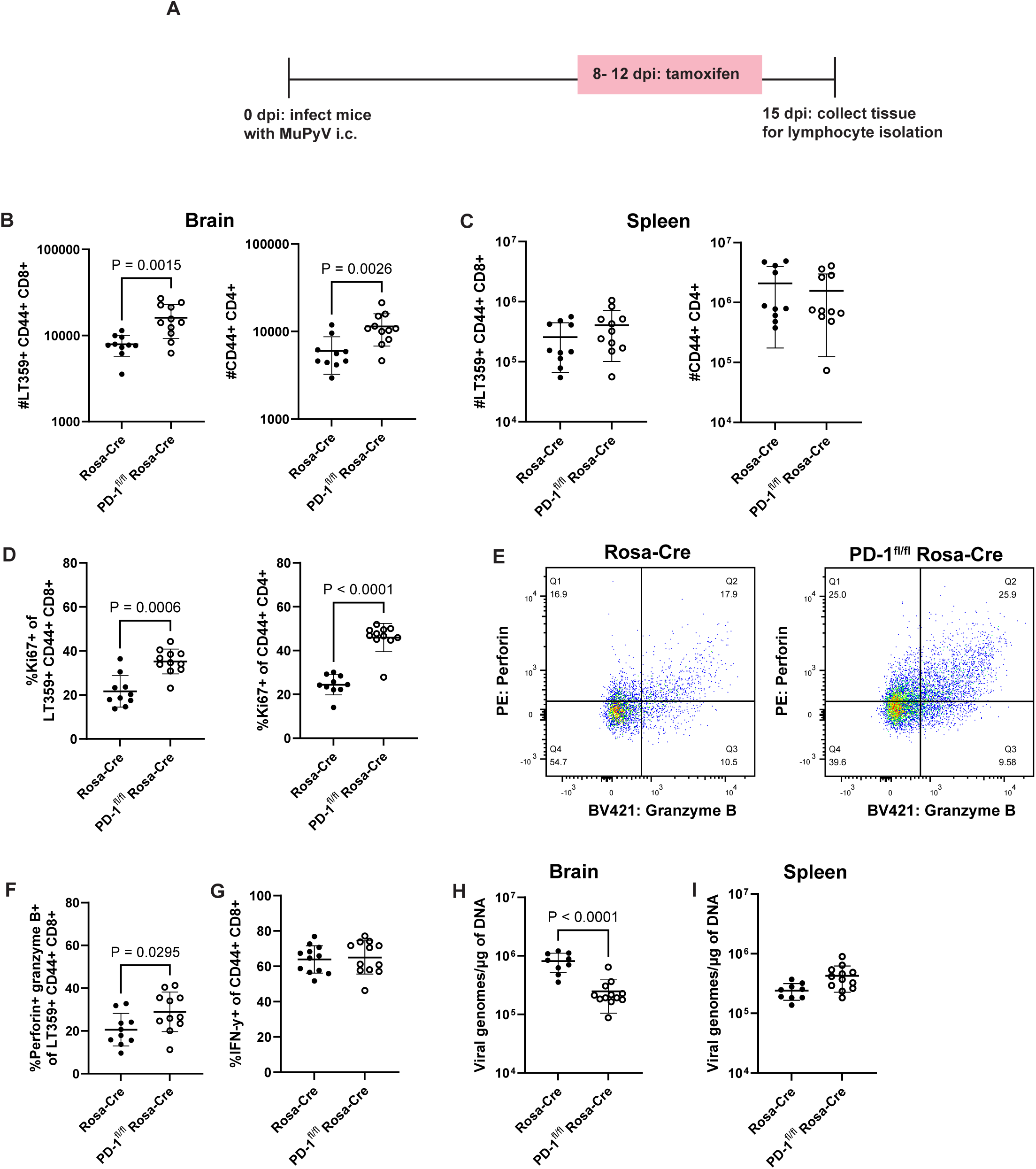
PD-1 alters the magnitude, function, and differentiation of brain T cells and limits viral control. PD-1^fl/fl^ Rosa-Cre and WT Rosa-Cre mice were treated as in A), which depicts the experimental scheme. B) Numbers of antigen-experienced CD4^+^ and virus-specific CD8^+^ T cells in the brains and C) spleens of Rosa-Cre and PD-1^fl/fl^ Rosa-Cre mice. D) Proportion of Ki67^+^ antigen-experienced CD4^+^ and virus-specific CD8^+^ T cells in Rosa-Cre and PD-1^fl/fl^ Rosa-Cre mice. E) Representative flow plots of perforin and granzyme B expression by virus-specific CD8^+^ T cells. Q2 indicates the proportion of cells that are double-positive for perforin and granzyme B, fully shown in F), which depicts percentages of double-positive perforin- and granzyme B-expressing virus-specific CD8^+^ T cells. G) Frequency of IFN-γ^+^ CD8^+^ T cells in Rosa-Cre and PD-1^fl/fl^ Rosa-Cre mice assessed through an *ex vivo* intracellular staining assay. H) Virus levels in brain tissue and I) spleen tissue, assessed through qPCR in in Rosa-Cre and PD-1^fl/fl^ Rosa-Cre. n = 10 Rosa-Cre mice and 11 PD-1^fl/fl^ Rosa-Cre mice. Error bars are mean ± SD. Data combined from two independent experiments. Statistical significance for graphical data calculated with a Mann-Whitney U test.

To ensure that these phenotypic changes were specific to brain-infiltrating T cells, we retro-orbitally injected mice with fluorophore-conjugated CD45 antibody immediately before euthanasia to mark intravenous (IV) or circulating cells^26^. With loss of PD-1, brain-infiltrating (i.e., IV^-^) T cells, showed increases in cell numbers, Ki67^+^ frequency, and perforin expression **(Supplementary 5A-C)**. Intravascular T cells (i.e., IV^+^) did not exhibit changes in cell numbers and Ki67^+^ frequency upon *Pdcd1* deletion **(Supplementary 5D-E)**. Although a statistically significant increase in perforin was detected, the degree of increase is more modest compared to the IV^-^ compartment **(Supplementary 5F**). Notably, induced *Pdcd1* deletion resulted in reduction in virus levels in the brain, but not in the spleen **(Fig. 2H-I).**

PD-1 deficiency can impair antibody responses by compromising the function of follicular helper T cells^20^. Sera from infected tamoxifen-treated Rosa-Cre and PD-1^fl/fl^ Rosa-Cre mice at 15 dpi showed no significant differences in levels of virus-specific IgG and IgM or IgG avidity against purified MuPyV virions (**Supplementary 6A-C**). In addition, CD4^+^ regulatory T (T_reg_) cells infiltrate brains of MuPyV infected mice^17^. A modest increase in the frequency and number of FOXP3^+^ CD25^+^ CD4^+^ T cells was seen in the PD-1^fl/fl^ Rosa-Cre mice **(Supplementary Fig. 7A-B)**.

We next asked whether PD-1 loss during late-stage persistent MuPyV CNS infection also led to T cell clonal expansion. Rosa-Cre and PD-1^fl/fl^ Rosa-Cre mice inoculated i.c. with MuPyV received tamoxifen 23 to 27 dpi, and T cells were isolated from brains at 50 dpi **(Fig. 3A)**.

**Figure 3.**
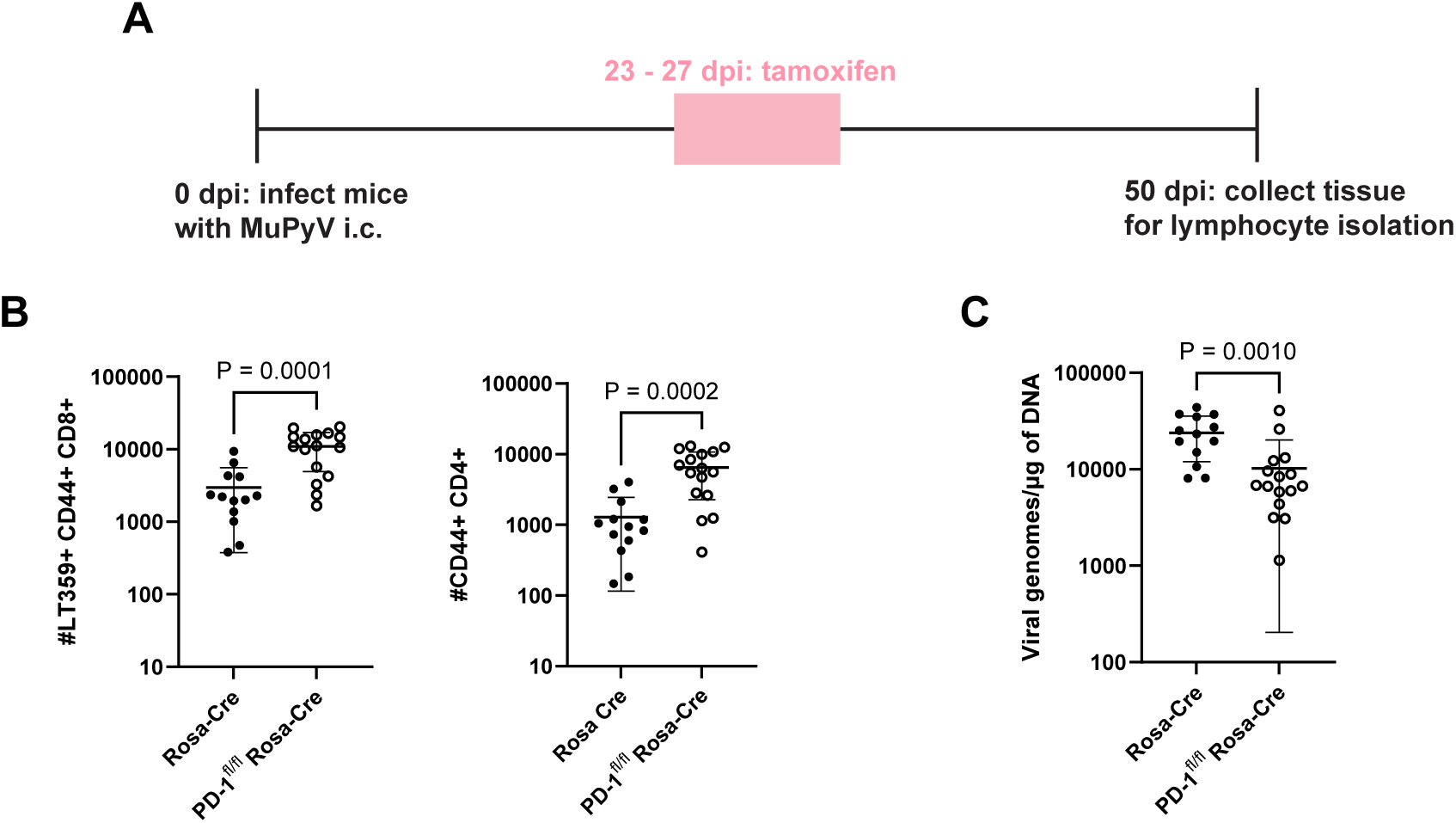
Loss of PD-1 during the late phase of the infection leads to expansion in T cells and reduced virus. A) Experimental setup. Rosa-Cre mice and PD-1^fl/fl^ Rosa-Cre mice were administered tamoxifen 23 to 27 dpi, and lymphocytes were isolated from brains at 50 dpi. B) Numbers of virus-specific CD8^+^ T cells and antigen-experienced CD4^+^ T cells. C) Viral genome copies. n = 13 Rosa-Cre mice and 16 PD-1^fl/fl^ Rosa-Cre mice. Data combined from two independent experiments. Statistical significance for B) and C) calculated with a Mann-Whitney U test.

Induced deletion of *Pdcd1* was associated with higher numbers of CD4^+^ and CD8^+^ T cell numbers along with reduced virus levels **(Fig. 3B-C)**. To determine if CD8^+^ T cell expansion with PD-1 deficiency was brain-autonomous, mice were depleted of circulating CD8^+^ T cells by systemic injection of anti-CD8b prior to tamoxifen administration 23 to 27 dpi, and T cells were harvested at 50 dpi **(Supplementary 8A)**. The anti-CD8b antibody does not bypass the blood-brain barrier. Thus, circulating CD8^+^ T cells were depleted in the blood **(Supplementary 8B)**, and brain-infiltrating CD8^+^ T cells were maintained **(Supplementary 8C)**. Under these conditions, CD8^+^ T cell numbers increased upon PD-1 loss, indicating that PD-1 limits expansion of brain-localized CD8^+^ T cells **(Supplementary 8D)**. Antigen-experienced CD4^+^ T cell numbers increased as well **(Supplementary 8E)**. Collectively, these data show that PD-1 loss results in brain-autonomous expansion of effector T cells and enhanced control of CNS MuPyV infection.

### Transcriptomic analysis of T cells from MuPyV-infected brains of WT and PD-1-deficient mice

To probe changes in the heterogeneity of the T cell response to MuPyV infection with loss of PD-1, we applied single-cell RNA sequencing (scRNA-seq) of CD4^+^ and CD8^+^ T cells from brains of PD-1^fl/fl^ Rosa-Cre and control Rosa-Cre mice. Both groups were injected i.c. with MuPyV, given tamoxifen 8 to 12 dpi, and lymphocytes isolated from brains at 15 dpi. CD44^hi^ CD4^+^ T and D^b^ LT359 tetramer^+^ CD8^+^ T cells were FACS-sorted from ten PD-1^fl/fl^ Rosa-Cre and ten Rosa-Cre mice and tagged with oligonucleotide bar-coded CD45 antibodies to distinguish strain and T cell type. CD4^+^ and CD8^+^ T cells were then respectively combined from PD-1^fl/fl^ Rosa-Cre and control Rosa-Cre mice and subjected to scRNA-seq.

Unsupervised clustering was performed and different clustering solutions were investigated before selecting a solution consisting of k = 11 clusters, as visualized in uniform manifold approximation and projection (UMAP) plots **(Fig. 4A)**. The eleven clusters were generated by t-distributed stochastic neighbor embedding (t-SNE) analysis, with four CD4^+^ T cell clusters, four CD8^+^ T cell clusters, and three clusters composed of both CD4^+^ and CD8^+^ T cells **(Fig. 4A)**. The CD4^+^ T cell clusters comprise a Th1-like effector population (cluster 2), regulatory T cells (cluster 4), a late effector population (cluster 5), and a stem-like population (cluster 6). Cluster 2 shows Th1-associated genes, such as *Tbx21* (Fang and Zhu, 2017) and *Ifng* (**Fig. 4B)**. Cluster 4 is strongly marked by Treg-associated genes, such as *Foxp3* and *Il2ra* **(Fig. 4C)**, and cluster 5 expresses *Cx3cr1*, which denotes highly differentiated effector CD4^+^ T cells^27,28^ (**Supplementary 9A)**. Notably, both clusters 5 and 2 express *Cxcr6,* a marker of effector CD4^+^ T cells^29,30^ **(Supplementary 9A)**. Cluster 6 shows expression of *Tcf7* (encoding TCF1, a transcription factor driving self-renewal of stem-like T cells^31^, *Slamf6*, and *Ccr7*, suggesting a stem-like phenotype for a subset of CD4^+^ T cells in the brains of MuPyV-infected mice^32^ **(Supplementary 9B**).

**Figure 4.**
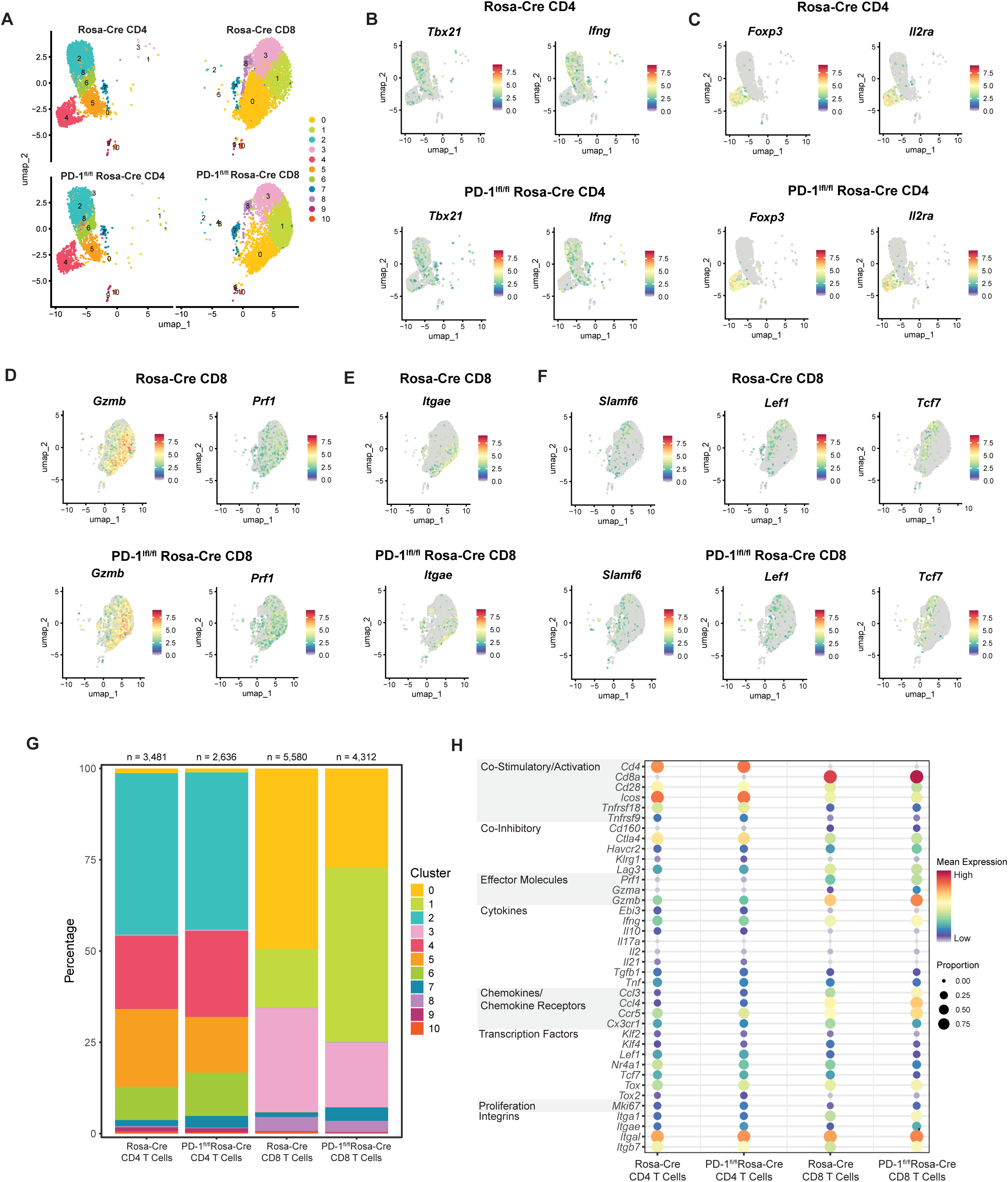
Single-cell RNA sequencing shows changes in resident-memory and effector CD8^+^ T cell clusters with loss of PD-1. Rosa-Cre and PD-1^fl/fl^ Rosa-Cre mice were infected and given tamoxifen 8 to 12 dpi. Virus-specific CD8^+^ T cells and antigen-experienced CD4^+^ T cells were sorted and processed for scRNA-seq analysis. A) UMAP showing clusters for each strain and cell type. B) Feature plots showing genes driving cluster 2 in the Rosa-Cre CD4^+^ T cell subset. C) Feature plots showing genes driving cluster 4 in the Rosa-Cre CD4^+^ T cell subset. D) and E) Feature plots showing genes driving cluster 1 in the Rosa-Cre CD8^+^ T cell subset. F) Feature plots showing genes driving cluster 3 in the Rosa-Cre CD8^+^ T cell subset. G) Bar plot demonstrating proportions of each cluster in the four groups. H) Dot plot showing expression (by mean expression and proportion of cells) of representative genes in the four groups. n = 10 mice per strain.

Similar to CD4^+^ T cells, CD8^+^ T cells show a cluster highly marked by *Cx3cr1* (cluster 0) **(Supplementary 9C)**. This overlaps with expression of *Irf8*, which has previously been implicated in effector CD8^+^ T cell differentiation^33^ **(Supplementary 9C)**. As cluster 0 also expresses *Prf1* and *Gzmb* (**Fig. 4D**), *Cx3cr1* expression suggests a late differentiated cytotoxic effector population^27^. Cluster 1 may be represented by a combination of effector and resident-memory precursors, as it also shows expression of *Gzmb* and *Prf1* along with expression of integrins associated with resident-memory, such as *Itgae* (**Fig. 4D-E**). Cluster 3 is marked by *Tcf7*, *Slamf6*, and *Lef1*, indicative of a stem-like or self-renewing phenotype^34,35^ **(Fig. 4F)**. Cluster 8 expresses *Malat1*, a long non-coding RNA that may regulate terminal effector cells^36^ (**Supplementary 9D**).

Compared to control Rosa-Cre CD8^+^ T cells, PD-1^fl/fl^ Rosa-Cre CD8^+^ T cells have a larger proportion attributed to cluster 1 and a lower proportion in cluster 0, suggesting that PD-1 regulates the balance between effector-like and resident-memory precursor CD8^+^ T cells and late differentiated effector CD8^+^ T cells **(Fig. 4G)**. This skewing toward effector differentiation by PD-1-deficient CD8^+^ T cells is supported by the increased expression of effector genes *Prf1*, *Gzma*, and *Gzmb* **(Fig. 4H and Supplementary 10A)**. PD-1^fl/fl^ Rosa-Cre CD8^+^ T cells also show increased expression of the integrin genes *Itgae* and *Itga1*, suggesting a more resident-memory-like phenotype **(Fig. 4H and Supplementary 10A)**. The proportion of PD-1^fl/fl^ Rosa-Cre CD8^+^ cells making up cluster 3 (stem-like cells) is decreased relative to control Rosa-Cre CD8^+^ T cells **(Fig. 4G)**. Reduced expression of *Tcf7*, *Lef1*, *Klf4*, and *Nr4a1*— genes associated with self-renewal of T cells^35,37,38^— further suggests that PD-1 loss results in a smaller subset of stem-like CD8^+^ T cells **(Fig. 4H and Supplementary 10A)**.

Changes in cluster proportions for the CD4^+^ T cells between Rosa-Cre and PD-1^fl/fl^ Rosa-Cre mice are more modest than for the CD8^+^ T cells, with a slight increase in the regulatory T cell cluster (cluster 4), a decrease in the *Cx3cr1* subset (cluster 5), and an increase in the *Tcf7* subset (cluster 6) **(Fig. 4G)**. For both CD4^+^ and CD8^+^ T cells, PD-1^fl/fl^ Rosa-Cre T cells have an increased proportion of cluster 7 **(Fig. 4G)**. Cluster 7 is marked by genes associated with proliferation and survival, such as *MKi67* (encoding Ki67)^39^, *Birc5*^40^, and *Stmn1*^41^**(Supplementary 9E-F)**. *MKi67* is increased in PD-1^fl/fl^ Rosa-Cre CD4^+^ and CD8^+^ T cells **(Fig. 4H and Supplementary 10B)**, signifying increased proliferation, as demonstrated in **Fig. 2D**. In summary, scRNA-seq analysis indicates an increase in effector-like and resident-memory-like CD8^+^ T cell subsets and decrease in stem-like CD8^+^ T cell subset with loss of PD-1 in the brains of MuPyV-infected mice.

### PD-1 deficiency shifts toward effector-like and resident-memory-like CD8^+^ T cell differentiation

We next sought to validate these T cell transcriptomic changes by flow cytometry using the same experimental setup (**Fig. 2A)**. Virus-specific CD8^+^ T cells were stained for TCF1, a marker for stem-like cell subsets encoded by *Tcf7*^31^, and CD103, a resident memory cell marker encoded by *Itgae*^42^. Comparing TCF1 and CD103 expression on virus-specific CD8^+^ T cells showed that brain-infiltrating CD8^+^ T cells do not co-express CD103 and TCF1 **(Fig. 5A).** PD-1 deficiency led to a higher fraction and number of CD103^+^ virus-specific CD8^+^ T cells **(Fig. 5B)**. Loss of PD-1 was also associated with a decrease in the frequency of anti-MuPyV CD8^+^ T cells expressing TCF1 **(Fig. 5C, left)**, but without a change in overall cell numbers **(Fig. 5C, right)**.

**Figure 5.**
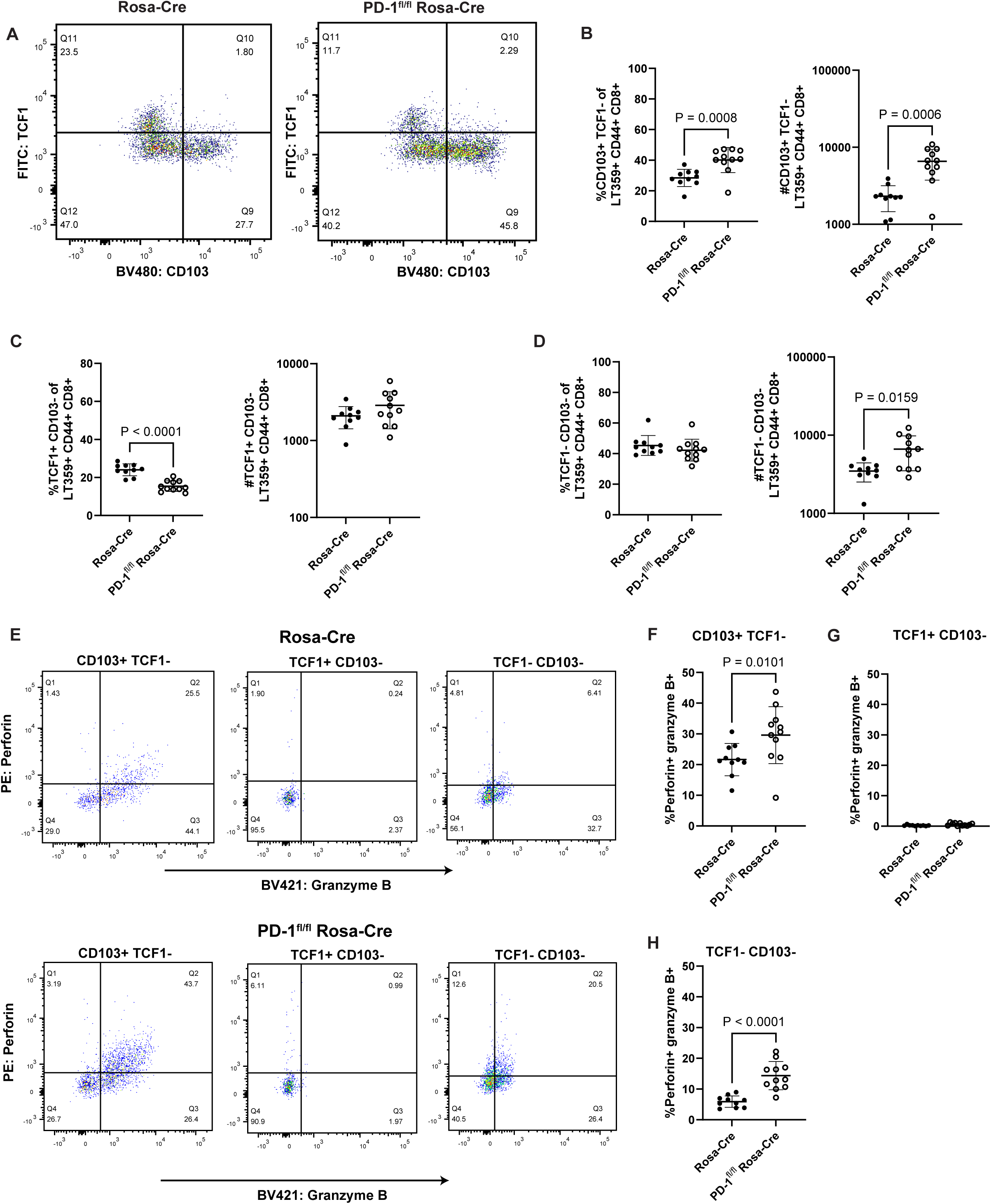
Loss of PD-1 skews differentiation of brain CD8^+^ T cells toward effector and resident-memory subsets. Rosa-Cre and PD-1^fl/fl^ Rosa-Cre mice were infected, given tamoxifen 8 to 12 dpi, and lymphocytes were isolated from the brain at 15 dpi. A) Representative flow plots showing virus-specific CD8^+^ T cells expressing CD103 and TCF1. B) Proportion and numbers of CD103^+^ virus-specific CD8^+^ T cells. C) Proportion and numbers of TCF1^+^ virus-specific CD8^+^ T cells. D) Proportion and numbers of CD103^-^ TCF1^-^ virus-specific CD8^+^ T cells. E) Representative flow plots of perforin- and granzyme B-expressing virus-specific CD8^+^ T cells in the three subsets shown in B, C, and D. F) Proportion of double-positive perforin- and granzyme B-expressing virus-specific CD8^+^ T cells that are TCF1^+^ CD103^-^. G) Same as F), but for TCF1^+^ CD103^-^ virus-specific CD8^+^ T cells. H) Same as F), but for double-negative TCF1^-^ CD103^-^ virus-specific CD8^+^ T cells. n = 10 Rosa-Cre mice and 11 PD-1^fl/fl^ Rosa-Cre mice. Error bars show mean ± SD. Data combined from two independent experiments. Statistical significance for all graphs was quantified with a Mann-Whitney U test.

Because T cell expansion with PD-1 loss occurs *in situ* in the brain **(Supplementary 8D-E)**, this result may indicate progression of TCF1^+^ CD8^+^ T cells into effector-competent and resident-memory cells. This possibility is supported by the finding that a higher number of double-negative CD103 and TCF1 CD8^+^ T cells was seen in the brains of PD-1 deficient mice without a change in frequency **(Fig. 5D)**. CD103 expression also marked the virus-specific CD8^+^ T cell subset with the highest proportion of cells positive for both perforin- and granzyme B, while TCF1 designated the cell subset with the lowest frequency of cells co-expressing these cytotoxic proteins **(Fig. 5E-H)**. With *Pdcd1* deletion, the TCF1^-^ cell subsets showed higher proportions of perforin- and granzyme B-expressing virus-specific CD8^+^ T cells **(Fig. 5F and 5H)**. Thus, transcriptomic and flow cytometric analysis together show that loss of PD-1 results in expansion of effector and resident memory-like CD8^+^ T cells.

### PD-1 loss intrinsic to brain CD8^+^ T cells does not drive T cell expansion or enhance antiviral control

Given the transcriptional changes in the PD-1-deficient CD8^+^ T cells and evidence that PD-1 intrinsically inhibits CD8^+^ T cell functionality^43,44^, we hypothesized that PD-1 signaling in CD8^+^ T cells per se limits their expansion and effector differentiation during MuPyV CNS infection.

We crossed PD-1^fl/fl^ mice to E8i-Cre^ERT2^ mice (E8i-Cre), generating a strain in which *Pdcd1* is deleted only in CD8^+^ T cells after tamoxifen administration (PD-1^fl/fl^ E8i-Cre). Using the same tamoxifen and infection regimen depicted in **Fig. 2A**, we assessed PD-1 expression on CD8^+^ and CD4^+^ T cells and verified that tamoxifen administration induced deletion of PD-1 in brain-infiltrating CD8^+^ T cells, but not CD4^+^ T cells **(Supplementary 11A-B)**. PD-1^fl/fl^ E8i-Cre mice and E8i-Cre control mice were infected with MuPyV i.c., given tamoxifen 8 to 12 dpi, and T cells isolated from the brain and spleen at 15 dpi. Negating our hypothesis, we found that loss of PD-1 only on CD8^+^ T cells resulted in no expansion, no change in frequency of Ki67^+^ and CD103^+^ cells, and no increase in perforin and granzyme B expression **(Fig. 6A-D)**. Moreover, virus levels in the brain were unchanged compared to infected E8i-cre control mice **(Fig. 6E)**.

**Figure 6.**
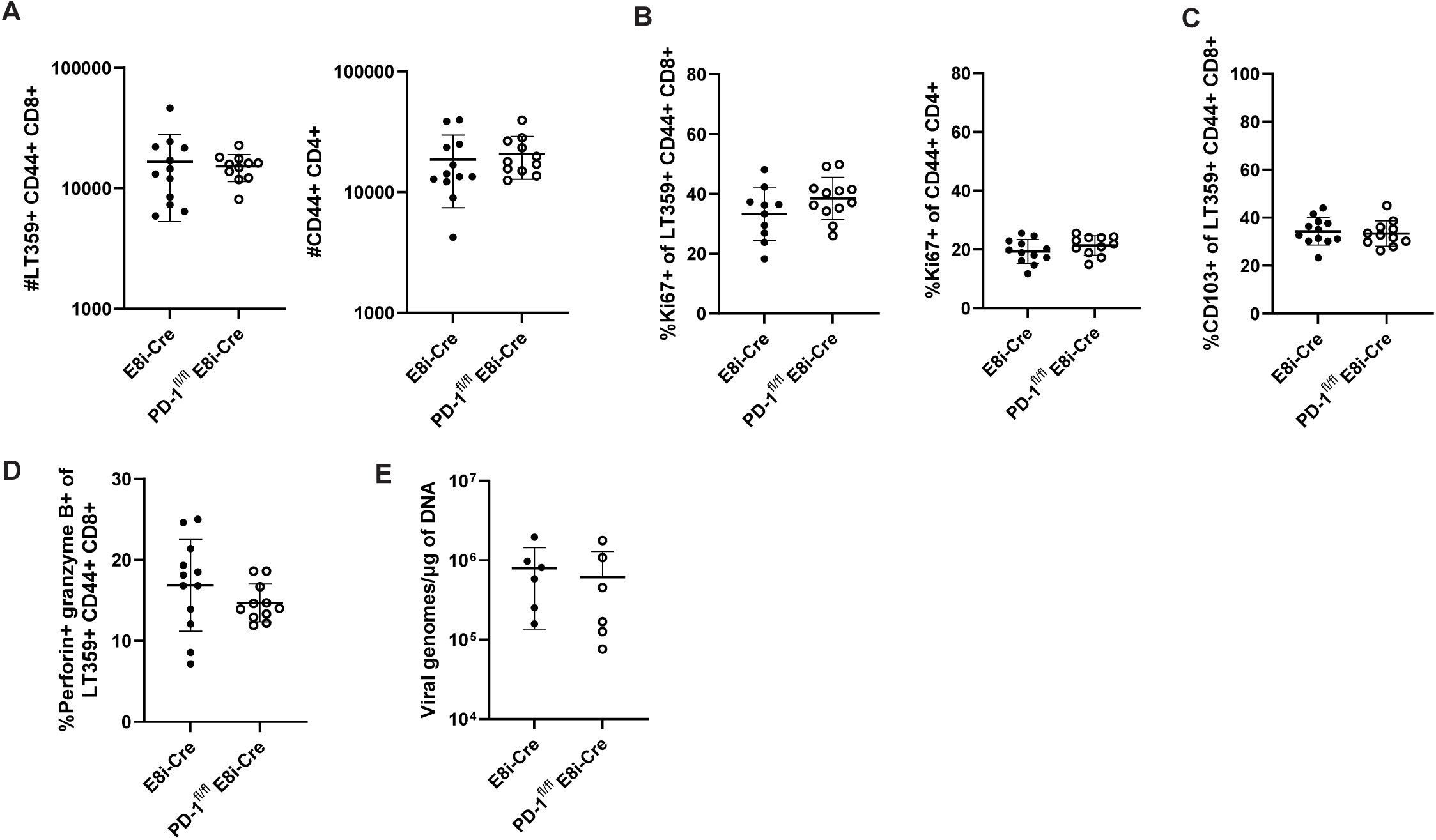
PD-1 loss intrinsic to CD8^+^ T cells does not drive their expansion and effector differentiation or enhance virus control. PD-1^fl/fl^ E8i-Cre and control E8i-Cre mice were infected and given tamoxifen 8 to 12 dpi. Lymphocytes were isolated from the brain at 15 dpi. A) T cell numbers. B) Proportion of Ki67^+^ of virus-specific CD8^+^ T cells and antigen-experienced CD4^+^ T cells. C) Proportion of CD103^+^ virus-specific CD8^+^ T cells. D) Proportion of perforin-and granzyme B-expressing virus-specific CD8^+^ T cells. E) Virus genome copies. n = 12 E8i-Cre and 11 PD-1^fl/fl^ E8i-Cre mice. Data from A) to D) are combined from two experiments, and E) shows data representative of two independent experiments. Error bars are mean ± SD. Statistical significance was calculated through a Mann-Whitney U test.

### PD-1 on brain-infiltrating CD4^+^ T cells inhibits CD4^+^ and CD8^+^ T cell proliferation and CD8^+^ T cell differentiation

Because CD4^+^ T cells in the MuPyV-infected brain also express PD-1, we next asked whether PD-1 loss intrinsic to CD4^+^ T cells would mimic the changes in CD8^+^ T cells seen in the MuPyV CNS infected, tamoxifen-treated PD-1^fl/fl^ Rosa-Cre mice. We bred PD-1^fl/fl^ mice to CD4-Cre^ERT2^ mice (CD4-Cre), generating PD-1^fl/fl^ CD4-Cre^ERT2^ mice (PD-1^fl/fl^ CD4-Cre), and confirmed that PD-1 is deleted on brain CD4^+^, but not CD8^+^, T cells after tamoxifen administration **(Supplementary 12A-B)**. PD-1^fl/fl^ CD4-Cre and CD4-Cre control mice were inoculated i.c. with MuPyV, given tamoxifen 8 to 12 dpi, and lymphocytes isolated from the brain at 15 dpi. Both CD4^+^ and CD8^+^ T cell counts were higher with CD4^+^ T cell intrinsic PD-1 deficiency, with an increase in the frequency of Ki67^+^ T cells in both populations **(Fig. 7A-B)**. Although the proportion of perforin- and granzyme B-positive virus-specific CD8^+^ T cells was not higher with loss of PD-1 on CD4^+^ T cells **(Fig. 7C)**, virus levels in the brains of PD-1^fl/fl^ CD4-Cre mice were reduced **(Fig. 7D)**. Similar changes in number and proportion as what was observed in the global PD-1 knockout mice **(Fig. 5B-D)** were seen in the PD-1^fl/fl^ Rosa-Cre mice. Virus-specific CD103^+^ CD8^+^ T cells expanded in proportion and number **(Fig. 7E)**, and TCF1^+^ CD8^+^ T cells were maintained with loss of PD-1 **(Fig. 7F)**. Double-negative effector TCF1^-^ CD103^-^ virus-specific CD8^+^ T cells expanded in number in the PD-1^fl/fl^ Rosa-Cre mice **(Fig. 7G)**. Similar to what was seen in the infected global PD-1 knockout mice, these findings demonstrate that PD-1 signaling on CD4^+^ T cells, not on CD8^+^ T cells, controls the magnitude and differentiation of virus-specific CD8^+^ T cells and the level of MuPyV infection in the brain.

**Figure 7.**
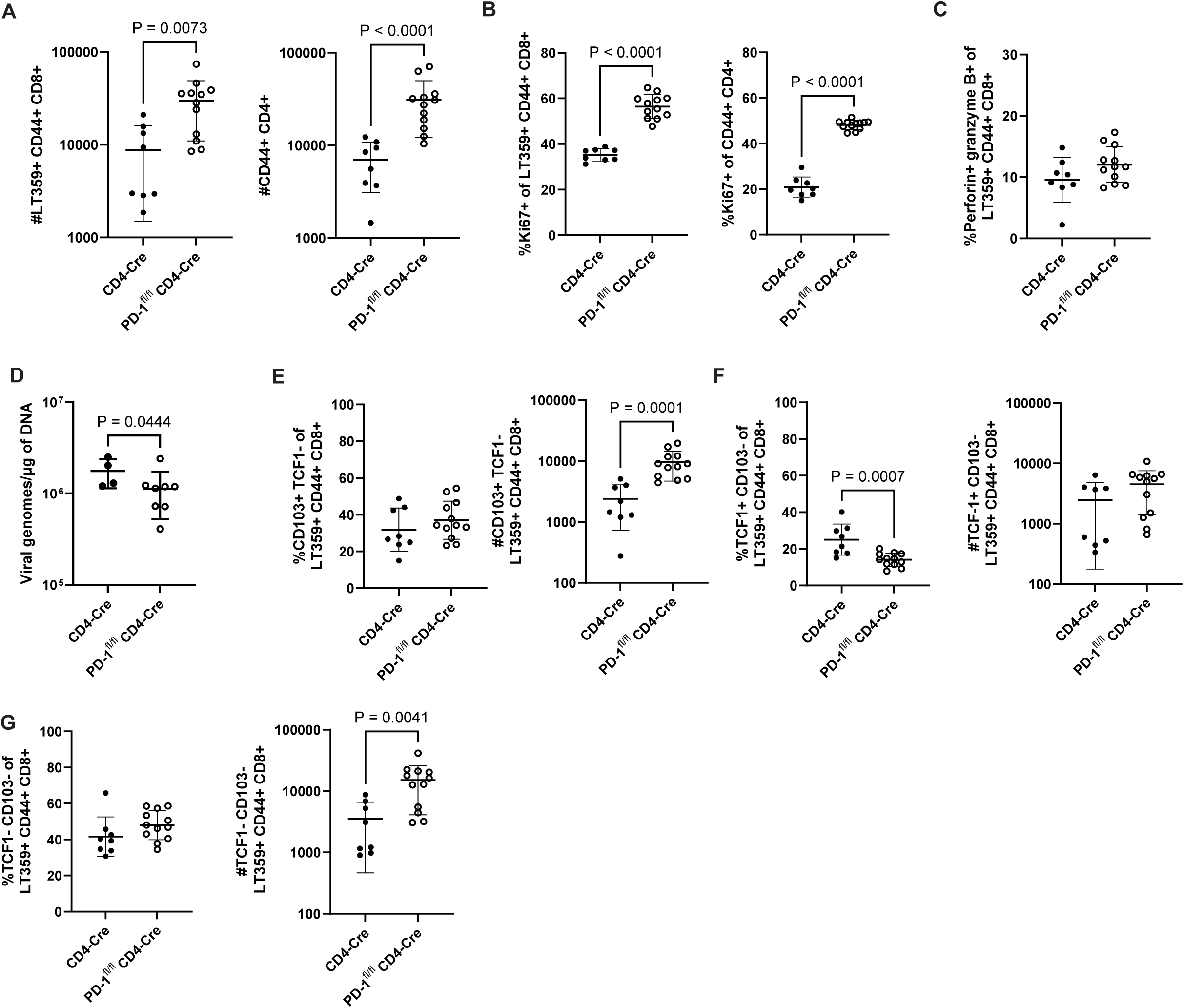
PD-1 on CD4^+^ T cells inhibits CD4^+^ and CD8^+^ T cell proliferation and CD8^+^ T cell differentiation. PD-1^fl/fl^ CD4-Cre and control CD4-Cre mice were infected, given tamoxifen 8 to 12 dpi, and at 15 dpi, lymphocytes were isolated from brain tissue. A) Numbers of virus-specific CD8^+^ T cells and antigen-experienced CD4^+^ T cells. B) Proportion of Ki67^+^ virus-specific CD8^+^ T cells and antigen-experienced CD4^+^ T cells. C) Proportion of virus-specific CD8^+^ T cells positive for both perforin and granzyme B. D) Virus levels in brain tissue. E) Frequency and number of CD103^+^ virus-specific CD8^+^ T cells. F) Frequency and number of TCF1^+^ virus-specific CD8^+^ T cells. G) Frequency and number of double-negative CD103^-^ TCF1-virus-specific CD8^+^ T cells. n = 8 CD4-Cre mice and 12 PD-1^fl/fl^ CD4-Cre mice. Error bars indicate mean ± SD. Statistical significance was calculated through a Mann-Whitney U test. Data from all graphs are combined from two independent experiments, except for D), which is representative of two experiments.

### PD-1 on CD4^+^ T cells regulates MuPyV-associated neuroinflammation

We next asked whether PD-1 on T cells in the brain controls MuPyV infection-associated neuroinflammation^17^. Microglia upregulate MHC class II (MHCII) expression in a neuroinflammatory microenvironment^45,46^. PD-1^fl/fl^ Rosa-Cre and Rosa-Cre mice were either sham or MuPyV injected i.c., then given tamoxifen 8 to 12 dpi. At 15 dpi, glial cells were isolated and phenotyped by flow cytometry, and microglia gated via CD45 and CD11b expression **(Fig. 8A)**. MHCII expression on microglia was higher in infected PD-1^fl/fl^ Rosa-Cre mice than infected Rosa-Cre mice **(Fig. 8B)**.

**Figure 8.**
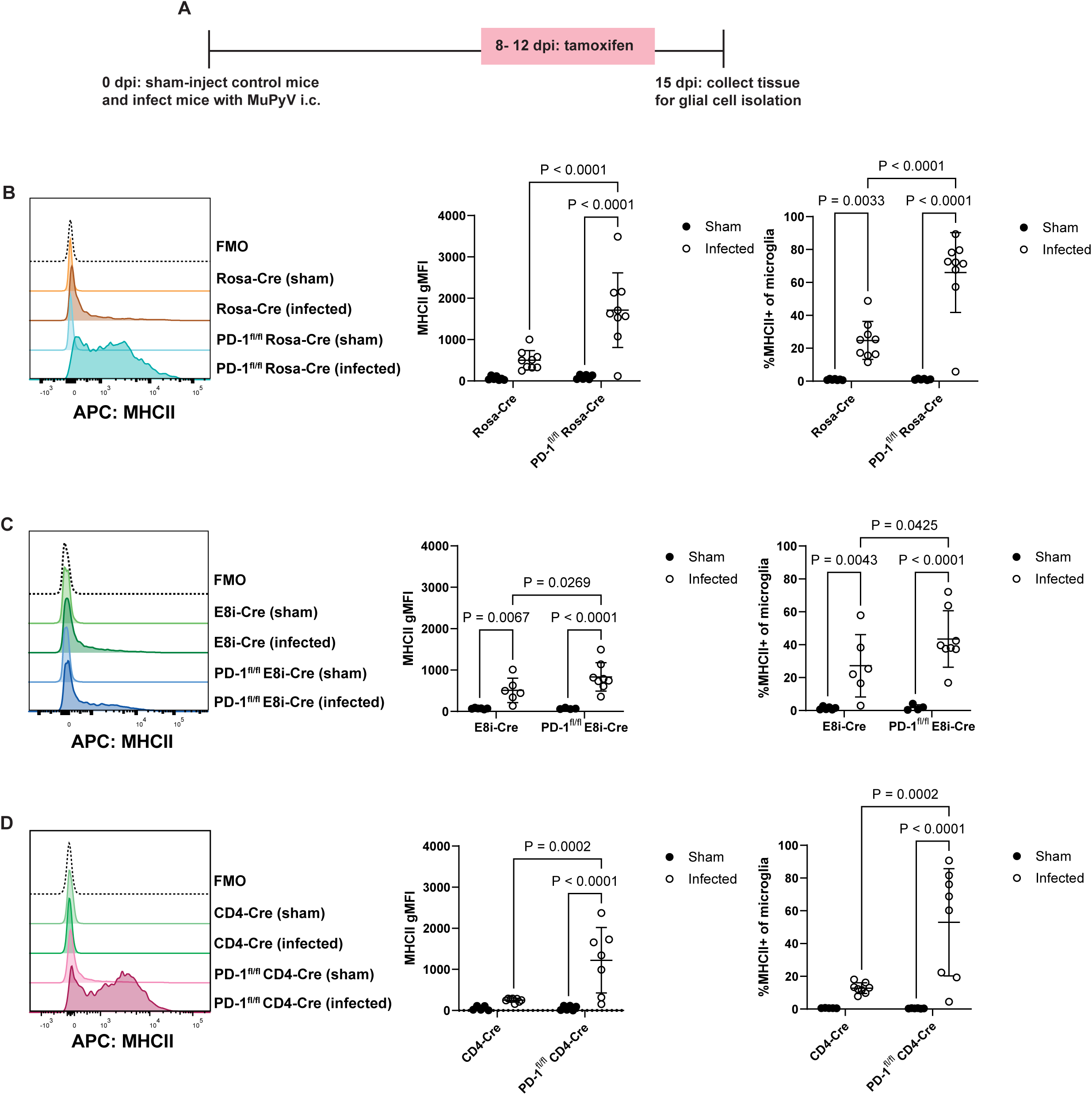
PD-1 on CD4^+^ T cells regulates MuPyV-associated neuroinflammation. Mice were sham- or MuPyV-injected i.c. At 8 to 12 dpi, mice were given tamoxifen, and at 15 dpi glial cells were isolated from brain tissue. A) MHCII expression in microglia of Rosa-Cre and PD-1^fl/fl^ Rosa-Cre mice. n = 7 Rosa-Cre mice (sham), 6 PD-1^fl/fl^ Rosa-Cre mice (sham), 10 Rosa-Cre mice (infected), and 9 PD-1^fl/fl^ Rosa-Cre mice (infected). B) MHCII expression in microglia of E8i-Cre and PD-1^fl/fl^ E8i-Cre mice. n = 6 E8i-Cre mice (sham), 5 PD-1^fl/fl^ E8i-Cre mice (sham), 6 E8i-Cre mice (infected), and 8 PD-1^fl/fl^ E8i-Cre mice (infected). C) MHCII expression in microglia of CD4-Cre and PD-1^fl/fl^ CD4-Cre mice. n = 5 CD4-Cre mice (sham), 6 PD-1^fl/fl^ CD4-Cre mice (sham), 8 CD4-Cre mice (sham), and 7 PD-1^fl/fl^ CD4-Cre mice (infected). Error bars are mean ± SD. Data combined from two independent experiments for each pair of strains. Statistical significance was calculated with a two-way ANOVA with uncorrected Fisher’s LSD post-test.

To determine the contribution of PD-1 on CD4^+^ and CD8^+^ T cells to MuPyV-associated neuroinflammation, glial cells were isolated from sham and infected mice with selective loss of PD-1 on CD4^+^ or CD8^+^ T cells. MHCII expression on microglia was only slightly increased in PD-1^fl/fl^ E8iCre mice than E8i-Cre mice **(Fig. 8C)**. *Pdcd1* deletion restricted to CD4^+^ T cells, however, resulted in increased microglia expression of MHCII (**Fig. 8D)** to levels matching those in the infected global PD-1 knockout mice **(Fig. 8B)**. These data suggest that PD-1 on CD4^+^ T cells, not on CD8^+^ T cells, counters neuroinflammation during MuPyV encephalitis.

## Discussion

PD-1 signaling on virus-specific T cells balances control of infection against tissue injury. PD-1-mediated restraint of CD8^+^ T cell function in non-lymphoid peripheral organs during persistent viral infections is well established. On the other hand, the impact of PD-1 inhibition of T cell activity in the setting of persistent viral encephalitis is incompletely understood. During polyomavirus infection in the CNS, CD4^+^ and CD8^+^ T cells both express PD-1. Whether PD-1 signaling operates equivalently or cooperatively on these T cell subsets to fine-tune their functionality is unknown. Here, we found that loss of PD-1 on CD4^+^ T cells is sufficient to promote expansion of effector, virus-specific CD8^+^ T cell in the brains of MuPyV-infected mice, resulting in improved viral control but at the expense of exacerbated neuroinflammation.

We employed a novel experimental setup to temporally delete *Pdcd1* during MuPyV CNS infection either globally or selectively on CD4^+^ or CD8^+^ T cells. Our previous studies used constitutive PD-L1 knockout mice and did not explore the contributions of PD-1 on individual T cell subsets^17,18^. Constitutive loss of PD-1 signaling may interfere with recruitment, differentiation, and expansion of virus-specific T cells^47,48^. Using the CD8-specific promotor E8i-Cre mice to selectively target CD8^+^ T cells for PD-1 loss on, Pauken et al. found that PD-1^-/-^ CD8^+^ T cells promoted antitumor immunity by PD-1-sufficient CD8^+^ T cells^6^. Here, we used a tamoxifen-inducible E8i-Cre system to delete *Pdcd1* after recruitment of T cells into the MuPyV-infected brain. Constitutive versus inducible PD-1 loss on CD8^+^ T cells may account for the apparent discrepancy between our data and this recent study as well as context-dependent differences in T cell behavior between the MuPyV-infected CNS and this tumor model.

Open questions from the present work include the nature of CD4^+^ T cell crosstalk with CD8^+^ T cells and how PD-1-deficient CD4^+^ T cells regulate PD-1-sufficient CD8^+^ T cells in the infected brain. Loss of PD-1 may alter CD4^+^ T cell cytokine profiles. First-line candidates are cytokines secreted by activated CD4^+^ T cells that act directly on neighboring CD8^+^ T cells. IFN-γ, secreted by Th1 CD4^+^ T cells, could augment viral epitope density and “titrate” PD-1 inhibition against increased TCR signaling. Another candidate cytokine is IL-21, critical to sustaining effector CD8^+^ T cell responses in chronic viral infections^7,29^. Our scRNA-seq results, however, failed to show upregulation of IFN-γ or IL-21 transcripts in PD-1-deficient CD4^+^ T cells **(Fig. 4H and Supplementary 10B)**. CD4^+^ T cells could also secrete cytokines that act indirectly via MHCII^+^ and/or MHCI^+^ viral antigen-presenting CNS-resident cells (e.g., microglia and astrocytes) or CNS-infiltrating myeloid cells. Crosstalk between microglia and T cells is documented but not well-understood. In the 5XFAD Alzheimer’s disease mouse model, CD4^+^ T cells can induce expansion of an MHCII^+^ subset of microglia associated with increased amyloid beta uptake and enhanced effector CD4^+^ Th1 cell function^49^.

Another possible mechanism by which PD-1 meditates CD4^+^-CD8^+^ T cell crosstalk is direct engagement of the PD-1/PD-L1 axis between CD4^+^ and CD8^+^ T cells. Brain-infiltrating T cells express PD-L1 **(Fig. 1B, Supplementary 1D, and Supplementary 2C)**. Because PD-1 loss only on CD8^+^ T cells does not alter CD8^+^ T cell function or numbers **(Fig. 6)**, PD-1 on CD8^+^ T cells may engage PD-L1 on CD4^+^ T cells to inhibit CD4^+^ T cell responses. Inhibitory “back-signaling” via PD-L1 on tumor-infiltrating CD4^+^ and CD8^+^ T cells has been reported^50^. PD-1 on T_reg_ cells in chronic LCMV infection can interact with PD-L1 on CD8^+^ T cells to enhance suppression of the antiviral CD8^+^ T cell response^51^. Similarly, a recent study demonstrated that PD-L1 on NK cells negatively regulates PD-1^+^ CD8^+^ T cells in the submandibular gland^52^. The possibility that PD-L1 engages PD-1 on CD8^+^ T cells (in *cis* or in *trans)* is excluded by the finding that PD-1 loss only on CD8^+^ T cells does not affect their expansion. Yet PD-1 loss on CD4^+^ T cells did not change perforin and granzyme B expression by CD8^+^ T cells (**Fig. 7C**), whereas expression of these effector molecules increased with global PD-1 deficiency (**Fig. 2E-F**), implying that PD-L1 back-signaling on CD8^+^ T cells may differentially affect effector activities. Notably, viral control was improved in both the global PD-1 knockout and the CD4-Cre-specific PD-1 knockout independently of changes in perforin and granzyme B expression (**Fig. 2H and Fig. 7D**), raising the question of how PD-1-deficient T cells contribute to control of neurovirulent polyomavirus infection. Virus-specific CD8^+^ T cells can control viral infection through non-cytolytic means^53^. Such mechanisms may be more likely during a neurovirulent infection given the nonrenewable nature of neurons, a key part of the CNS^54^.

PD-1 impacts the differentiation of CD8^+^ T cells^31,47^. During MuPyV encephalitis, global loss of PD-1 increased TCF1^-^ effector-like and resident-memory-like CD103^+^ CD8^+^ T cell numbers, while maintaining a proliferation-competent TCF1^+^ CD8^+^ T cell population **(Fig. 5).** Virus-specific CD8^+^ T cells showed mutually exclusive expression of CD103 and TCF1. This recapitulates a previous observation that virus-specific CD103^+^ CD8^+^ T cells in B6 mice infected with MuPyV i.c. exhibited lower TCF1 expression compared to CD103^-^ CD8^+^ T cells in the same mice^18^. Although TCF1 marks T cells positioned for a memory trajectory, TCF1 also identifies precursor-exhausted T cells. Precursor-exhausted T cells are generated during both acute and chronic infections; self-renewal of these TCF1^+^ cells by asymmetric division replenishes the short-lived effector T cells that are responsive to PD-1 blockade during chronic LCMV infection^31,55,56^. An intriguing possibility is that TCF1^+^ CD8^+^ T cells are similarly maintained during persistent CNS viral infections.

CD103, an αE integrin that complexes with b7 to bind epithelial cadherin, is a commonly used marker of resident-memory CD8^+^ T (T_RM_) cells^42^. CD103 is expressed on a fraction of MuPyV-specific brain T_RM_ cells, with CD103^+^ brain T_RM_ cells exhibiting increased proliferation upon viral rechallenge^17,18^. The increase in the proportion of virus-specific CD103^+^ CD8^+^ T cells after loss of PD-1 **(Fig. 5B)** parallels the increase in brain-infiltrating CD103^+^ CD8^+^ T cells seen in MuPyV-infected PD-L1^-/-^ mice^17^. Preferential expansion of CD103-expressing PD-1^-/-^ CD8^+^ T cells has been described in other CNS viral infection mouse models^57,58^. Analyzing the endogenous, polyclonal CD8^+^ T cell response with *Pdcd1* deletion induced after T cell entry into the infected CNS clearly departs from most studies that infer a PD-1-intrinsic effect based on the fate and function of adoptively transferred PD-1^-/-^ TCR-transgenic CD8^+^ T cells^43,44,47^. Our work highlights the importance of using mouse models that approximate physiological contexts to investigate mechanisms of PD-1 immunoregulation *in vivo*.

PD-1 immunotherapy has a checkered track record in mitigating PML morbidity and mortality. The pathogenesis of neurological irAEs post-ICI therapy is incompletely understood, but may involve overactivation of microglia^59^. Here, we assessed neuroinflammation by MHCII expression on microglia and demonstrated that PD-1 loss on CD4^+^ T cells inhibited MHCII expression to a similar degree as global PD-1 loss (**Fig. 8A and 8C**), suggesting that PD-1 on CD4^+^ T cells is critical to restraining MHCII-associated neuroinflammation. These results align with recent evidence of an association between clonally expanded effector CD4^+^ T lymphocytes and neurological irAEs with ICI therapy^60^. Our results imply that CD4^+^ T cells are necessary for an effective response to ICI in the CNS, raising the central conundrum: how to balance injurious neuroinflammation against PD-1 blockade-mediated viral control. In this connection, spleen tyrosine kinase (Syk) was recently identified as a target to reduce microglial activation and neuroinflammation^61^.

Retrospective analysis of PML patients with inherited and acquired T cell deficiencies point toward low CD4^+^ T cell counts as a central risk factor for PML^62^. Association between particular HLA class II alleles and protection against JCPyV, and identification of JCPyV mutations in CD4^+^ T cell epitopes, further support the importance of CD4^+^ T cell immunity for controlling JCPyV infection^63,64^. Additionally, multiple sclerosis patients treated with natalizumab, an anti-α4 integrin biologic associated with PML^65^, have reduced CD4^+^:CD8^+^ T cell in the CSF but normal ratios in peripheral blood^66^. Evidence presented here reveals that PD-1 on CD4^+^ T cells is key to modulating both antiviral CD8^+^ T cell responses and neuroinflammation in the CNS. These data further suggest that CNS-infiltrating CD4^+^ T cells should be considered when stratifying PML patients for benefit from PD-1 checkpoint therapy.

## Methods

### Mice

Rosa26-Cre^ERT2^ mice (Rosa-Cre) were purchased from The Jackson Laboratory (strain #004847) and bred to PD-1^fl/fl^ mice. PD-1^fl/fl^ mice were also bred to E8i-Cre^ERT2^ (E8i-Cre) mice, provided by Dario Vignali at the University of Pittsburgh^67^, and CD4-Cre^ERT2^ (CD4-Cre) mice, purchased from The Jackson Laboratory (strain #022356). Mice were bred and housed in pathogen-free facilities. Both male and female mice were used for experiments at 6 to 12 weeks of age. All mouse experiments were approved by the Penn State College of Medicine Institutional Animal Care and Use committee (Protocol PRAMS201447619).

### Virus injection and tamoxifen administration

At 6 to 12 weeks old, mice were inoculated i.c. with MuPyV, strain A2, in Dulbecco’s Modified Eagle’s Medium (DMEM) with 5% FBS. Inoculation was performed by injecting the anatomical right frontal lobe with 30 μl of 1 × 10^7^ plaque-forming units (PFU) of virus, as previously described^21^. Sham injections were performed similarly with vehicle or plain DMEM. For tamoxifen administration, tamoxifen (Sigma) was prepared at a concentration of 20 mg/ml in corn oil (Sigma) and mixed at 37°C for 3 h. Except for the CD4-Cre mice, all adult mice (weighing approximately 20 g) with inducible Cre^ERT2^ were given 100 µl (2 mg) of tamoxifen i.p. consecutively for 5 days. As we found a higher dose induced better recombination for the CD4-Cre mice, both strains (CD4-Cre and PD-1^fl/fl^ CD4-Cre) were given 3 mg of tamoxifen using the same dosing schedule. For T cell depletions, mice were injected i.p. once per week with 250 μg of anti-CD8b (clone H35-17.2) or control rat IgG (Jackson Immunoresearch) prepared in phosphate-buffered saline (PBS).

### Generation of PD-1^fl/fl^ mice

Two single-guide RNAs (sgRNAs) were generated to target *Pdcd1* 5’ upstream of exon 2 in the surrounding intron: 64_Pdcd1_sgRNAup1: AGGAGCTTGTAGCTTCTTGT and 65_Pdcd1_sgRNAup2: GAAGCTACAAGCTCCTAGGT. Additionally, two sgRNAs were generated to target 3’ downstream of exon 2 in the surrounding intron: 66_Pdcd1_sgRNAdw1: GGCATCTGAGGATTTCCACA and 67_Pdcd1_sgRNAdw2: GGGCATCTGAGGATTTCCAC.

sgRNAs were selected using the CRISPR Guide RNA design tool Benchling (Benchling). Additionally, a donor plasmid containing the desired LoxP sequences and homology arms surrounding exon 2 was introduced for homology-directed repair. Cas9/plasmid injection was performed on C57BL/6NJ zygotes. Founders were genotyped via long-range PCR and sequencing at the Pdcd1 locus. Long-range PCR primers of PD-1-/- mice: 76_Pdcd1_upF1: TCCCACTGACCCTTCAGACAG and 79_Pdcd1_RHdR2: CGTGTCAGGCACTGAAGAGATC. Products were subsequently sequenced with 77_Pdcd1_upF2: AACTAGGCTAGCCAACCAGAAG and 81_Pdcd1_seqF: ACAGTGGCATCTACCTCTGTGG. The resulting mouse line was backcrossed to C57BL/6J (Jackson Laboratory, strain #000664).

### Cell isolation from tissues

For IV labeling, mice were anesthetized by isofluorane inhalation and injected retro-orbitally with 3 μg fluorophore-conjugated (PerCPCy5.5) CD45 antibody (Biolegend)^26^. After 3 min, mice were euthanized. For lymphocyte isolation from brain tissue, brains were minced, digested with collagenase type I (Worthington) and deoxyribonuclease I (DNAse) (Worthington) for 20 min at 37°C, then passed through a 70 µm filter. Lymphocytes were then separated via a 44%/66% Percoll (Cytiva) step gradient, collected from the interface, and resuspended in wash media [DMEM with 2% fetal bovine serum (FBS)]. For lymphocyte isolation from spleens, spleens were passed through a 70 µm filter, and red blood cells (RBC) were lysed with ammonium-chloride-potassium (ACK) buffer for 5 min at room temperature (RT). Single cells were then resuspended in wash media.

For glial cell isolation, brains were dissected, minced, and digested with collagenase type I (Worthington) for 20 min at 37°C. Homogenate was passed through a 100 µm filter, centrifuged on a 37% Percoll gradient, then single cells were resuspended in wash media.

For blood samples, blood was collected from mice through submandibular bleeds into tubes containing an equal volume of heparin. RBCs were lysed with two 5-min ACK treatments, washed with wash media, and resuspended in 500 µl of wash media.

### Intracellular staining assay

Cells were incubated in a 96-well round-bottom plate (Corning Costar) in DMEM with 10% FBS containing brefeldin A (BD Biosciences) with either LT359-368 peptide (SAVKNYAbuSKL) (1 μM)^68^ or no peptide for five hours, after which cells were stained with antibodies.

### Antibody staining and flow cytometry

Cells were transferred to a 96-well round-bottom plate (Corning Costar), stained with a fixable viability dye (Thermo Fisher Scientific) and Fc block (BD Biosciences) for 20 min at 4°C, stained with fluorophore-conjugated antibodies to cell surface molecules (see **Supplementary Table 1** for antibody list) for 30 min at 4°C, fixed with 4% PFA for 15 min at RT, then resuspended in FACS buffer [PBS with 1% bovine albumin (Thermo Fisher Scientific) and 0.01% NaN_3_] For intracellular antibody staining, cells were treated with permeabilization/fixation buffer from the eBioscience™ Foxp3 / Transcription Factor Buffer Set (Thermo Fisher Scientific) for 15 min at RT, then incubated with fluorophore-conjugated antibodies (**Supplementary Table 1**) for 30 min at RT. Samples were then washed and resuspended in FACS buffer. Flow cytometry data acquisition was performed on a 23-color BD FACS Symphony flow cytometer (BD Biosciences) and analyzed with FlowJo (BD Life Sciences). To minimize spillover between colors in multicolor panels, compensation controls for each utilized color were used. Fluorescence minus one (FMO) controls were also used as necessary.

### Apoptosis assay

After cell isolation, cells were resuspended in PBS and stained with a fixable viability dye (Thermo Fisher Scientific) for 20 min at 4°C. Cells were then washed and stained for surface antibodies and tetramer in PBS for 30 min at 4°C, after which cells were resuspended in 1X annexin binding buffer (BD Biosciences) containing annexin V (Biolegend). Cells were incubated for 15 min at room temperature in the dark, then resuspended in binding buffer and analyzed by flow cytometry.

### Quantitative PCR for MuPyV genomes

DNA was extracted from tissue using the Wizard Genomic DNA Purification Kit (Promega). Viral DNA was then quantified by Taqman qPCR and viral genome copy numbers determined as described^68^.

### Serum collection, ELISA, and avidity assay

Blood samples were collected via submandibular bleeding and allowed to clot at room temperature for thirty min, then spun down at 2000 x g. Serum was collected into fresh tubes and frozen in aliquots at −80°C. IgG concentration and IgM detection were performed using EIA/RIA Polystyrene High Bind Microplate (Corning Incorporated) coated overnight at 4°C with 1×10^6^ genome/well of Opti-prep purified MuPyV. Plates were blocked with 1% BSA in 0.1% Tween PBS (blocking buffer). Mouse serum was diluted 1:150 in blocking buffer before being added to the plate. For the avidity assay, the plates were treated the same way as the ELISA, but virus:Ab complexes were treated with 2M NH_4_SCN in 0.1M phosphate for 15 min. Bound IgG in the ELISA and avidity assay was detected with an anti-mouse IgG specific secondary conjugated with horseradish peroxidase (HRP) (Bethyl Laboratories). IgM was detected using an anti-mouse IgM specific secondary Ab conjugated with biotin (BioLegend) and streptavidin bound to HRP (BioLegend). Plates were developed with 1-Step Ultra TMB-ELISA (Thermo Fisher Scientific) and imaged using the Synergy HI plate reader with absorption at O.D. 450. IgG concentration was calculated using a standard curve of the VP1-specific rat mAb 8H7A5, which was detected with an anti-rat IgG specific secondary Ab conjugated with HRP (BioLegend). IgG concentrations in the avidity assay at 2M NH_4_SCN were normalized to the absorption of the same sample treated with 0M NH_4_SCN in 0.1M phosphate.

### Tissue preparation for MERFISH transcriptomics and analysis

Brains were cut on the sagittal plane and prepared as previously described^22^. MERFISH analysis was also performed as previously described^22^.

### Preparing samples for BD Rhapsody^TM^ single cell capture system

10 Rosa-Cre and 10 PD-1^lf/fl^ Rosa-Cre mice were infected with MuPyV (A2 strain) and given tamoxifen 8 to 12 days dpi. Mice were IV-labeled, brains resected, and lymphocytes isolated and stained with CD3, CD8, CD4, CD44 (all Biolegend), and D^b^ LT359 tetramers (NIH Tetramer Core). After staining, cells were pooled per mouse strain and stored overnight in BD OMICS-Guard Sample Preservation Buffer (BD Biosciences). The next day, cells were sorted into CD3^+^ CD4^+^ CD44^+^ T cells and CD3^+^ CD8^+^ CD44^+^ D^b^ LT359^+^ T cells using a BD FACSAria SORP (BD Biosciences). After sorting, cells were multiplexed by cell type and strain using the BD Ms Single-Cell Multiplexing Kit (BD Biosciences) per manufacturer’s instructions. Cells were loaded onto the BD Rhapsody^TM^ Cartridge, and mRNA was hybridized onto BD Rhapsody™ Enhanced Cell Capture Beads. Beads were retrieved, and reverse transcription was performed as detailed in the BD Rhapsody™ Single-Cell Analysis System User Guide.

### Library preparation and whole transcriptome sequencing

Libraries were prepared in the Penn State College of Medicine Genome Sciences core using BD Rhapsody™ System mRNA Whole Transcriptome Analysis (WTA) and Sample Tag following the manufacturer’s protocol with the suggested cycles of 12 and 11 for RPE PCR and Sample Tag PCR1, respectively. For indexing, 8 cycles for WTA and 6 for cycles for sample indexing were carried out. Final libraries were assessed for size distribution and concentration using BioAnalyzer High Sensitivity DNA Kit (Agilent Technologies). The libraries were pooled and sequenced on Illumina NovaSeq 6000 (Illumina) to get on average of 35,000 reads per cell, paired end 71 bp (read 2) and 51 bp (read 1) reads, according to the manufacturer’s instructions. Samples were demultiplexed using BCL Convert software (Illumina). Adaptors were not trimmed during demultiplexing.

### scRNA-seq whole transcriptome analysis

The Seurat R package v. 5.3.0^69^ was used to analyze a Seurat data object produced by the BD Rhapsody^TM^ pipeline. Quality control procedures were applied to filter low quality cells, and these were based on data-driven thresholds for the number of genes detected in each cell, the number of molecules per cell, and mitochondrial gene percentages. A total of 16009 cells were identified that were amenable for analysis. Read counts were normalized using Seurat’s “LogNormalize” procedure, then the “vst” method was applied to perform feature selection.

After scaling the data, dimension reduction was performed using the “pca” approach. A nearest-neighbor graph was created, then clustering was conducted with different values of the “resolution” parameter in Seurat’s FindClusters function. Exploratory analyses, including silhouette plots, dendrograms, and gene expression heatmaps of differentially expressed genes were used to identify an optimal clustering solution. Expression patterns for select genes of interest were visualized in feature plots based on UMAP embeddings. Dot plots were created to display proportions of cells expressing genes of interest as well as mean expression levels. All analyses were performed with R v. 4.4.3 (R Core Team 2025) software. Feature plots were generated with Seurat; custom R scripts were used to generate all other figures, some of which leveraged the ggplot2 R package (RRID:SCR_014601). The dplyr (RRID:SCR_016708) and tidyr (RRID:SCR_017102) R packages were used to perform data manipulations.

### Statistical analysis

Flow cytometry data was analyzed by FlowJo (BD Biosciences). Graphs were generated and statistical tests performed using GraphPad Prism (Dotmatics). Statistical tests are detailed in each figure legend. In general, datasets with two independent groups were analyzed with a Mann-Whitney U test. Datasets with more than two dependent groups were analyzed with Friedman’s test, a non-parametric version of repeated measures one-way ANOVA, and Dunn’s post-test.

Datasets comparing two independent variables were analyzed with a two-way ANOVA with uncorrected Fisher’s LSD post-test. Graphical data is shown as datapoints with mean and error bars indicating standard deviation. p values <0.05 are considered significant; all statistically significant p values are shown.

## Author contributions

Writing and editing: AB and AEL

Experimental design and execution: AB, AEL, SAS, MA, AP, KMA, KNA, TDS, MDL, GJ, SMB, LB, and JAA

Data analysis: AB, EA, and VW

## Acknowledgments

We would like to acknowledge all members of the Lukacher lab for their feedback on this manuscript. We would also like to acknowledge the Genomics Core Facility (RRID:SCR_021123), the Flow Cytometry Core (RRID: SCR_021134), and the Department of Comparative Medicine at the Penn State College of Medicine for their respective services and assistance.

