## Supplemental Figs 1-2 for "PD-1 regulates CD4^+^ T cell-mediated CD8^+^ T cell responses in the brain to balance viral control and neuroinflammation"

### Supplementary 1

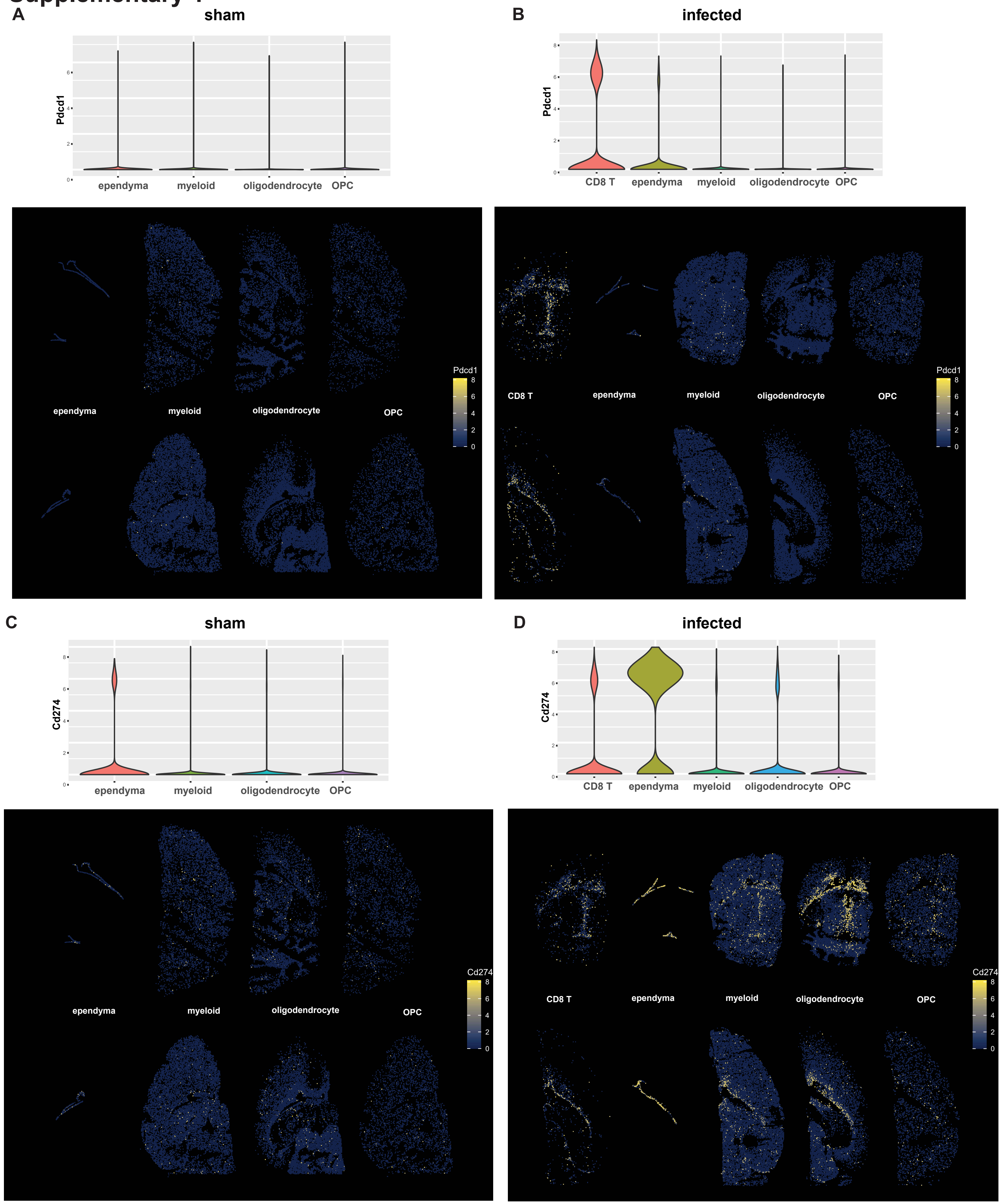


**Supplementary 1. MERFISH reveals spatial expression of Pdcd1 and Cd274 in the MuPyV-infected brain.** WT mice were sham-injected with vehicle or infected with MuPyV i.c. Brains were collected at 8 dpi. **A)** Violin plot of *Pdcd1* expression in multiple cell types, along with two representative brain sections, in sham-injected mice. Spots indicate 8 x 8 µm spots, as determined by an image-based segmentation method[22], and are positioned according to their corresponding x and y coordinates, regardless of z coordinates. Spots are colored according to log-normalized *Pdcd1* transcript expression. **B)** shows *Pdcd1* expression similarly to A) in MuPyV-infected mice. **C)** shows *Cd274* expression in sham-injected mice, while **D)** shows *Cd274* expression in MuPyV-infected mice. n = 2 mice per condition.

### Supplementary 2

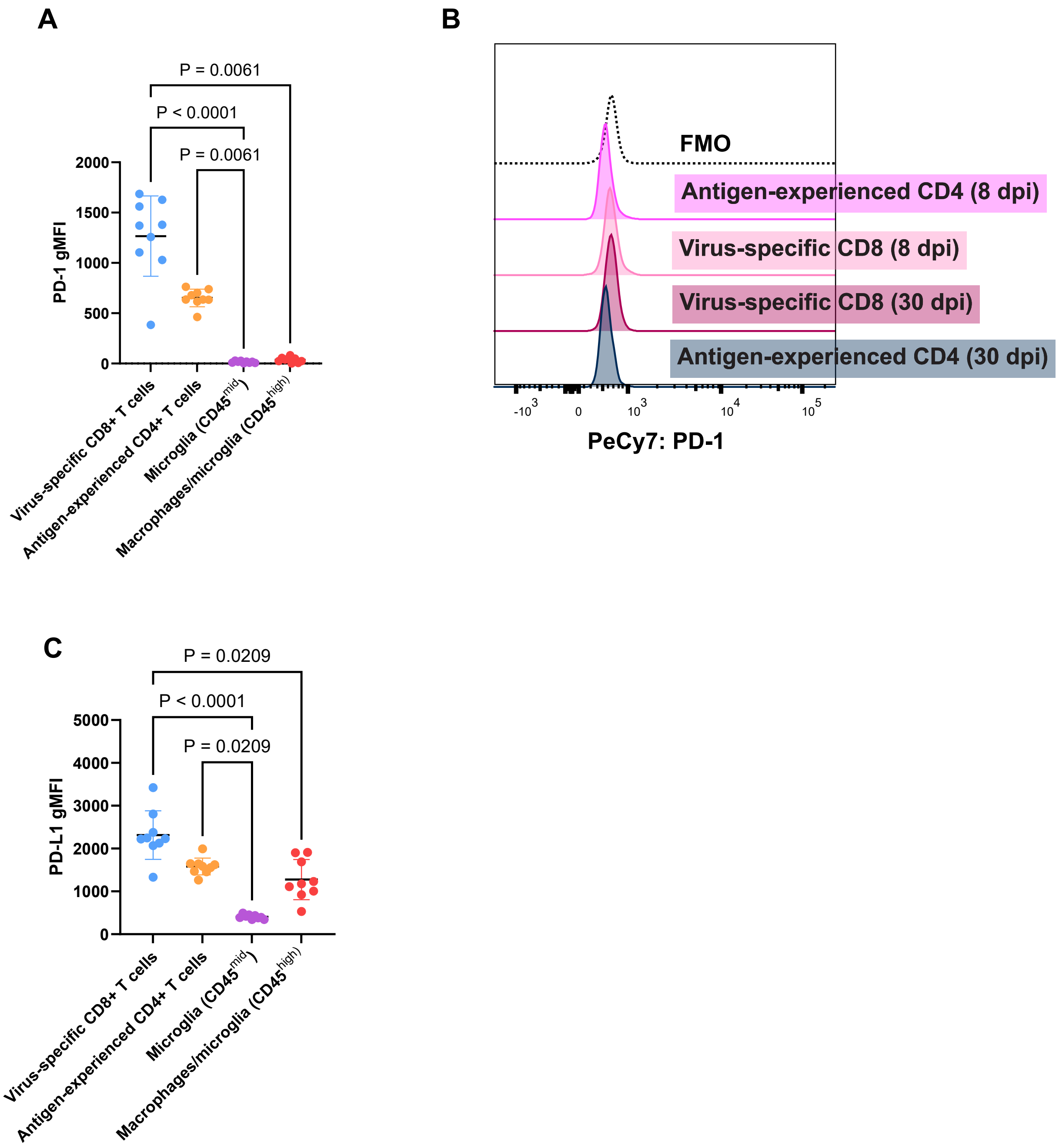


**Supplementary 2. PD-1 and PD-L1 expression are maintained on brain-infiltrating T cells during persistent infection.** WT mice were infected with MuPyV i.c., and at 8 and 30 dpi. Cells were isolated from the brain and blood and analyzed for PD-1 and PD-L1 expression. Gating strategies are specified in Figure 1. **A)** PD-1 expression on different cell types in the brain at 30 dpi. **B)** Representative histogram of PD-1 expression on T cells in the blood at 8 and 30 dpi. **C)** PD-L1 expression on different cell types in the brain at 30 dpi. n = 8 mice for 8 dpi and 9 mice for 30 dpi. Error bars indicate mean ± SD. Data are representative of two independent experiments. As each mouse yielded all the immune cell types shown, statistical significance for A) and C) was calculated with the Friedman's test, a nonparametric alternative to the repeated measures one-way ANOVA, with a Dunn's post-test.

### Supplementary 3

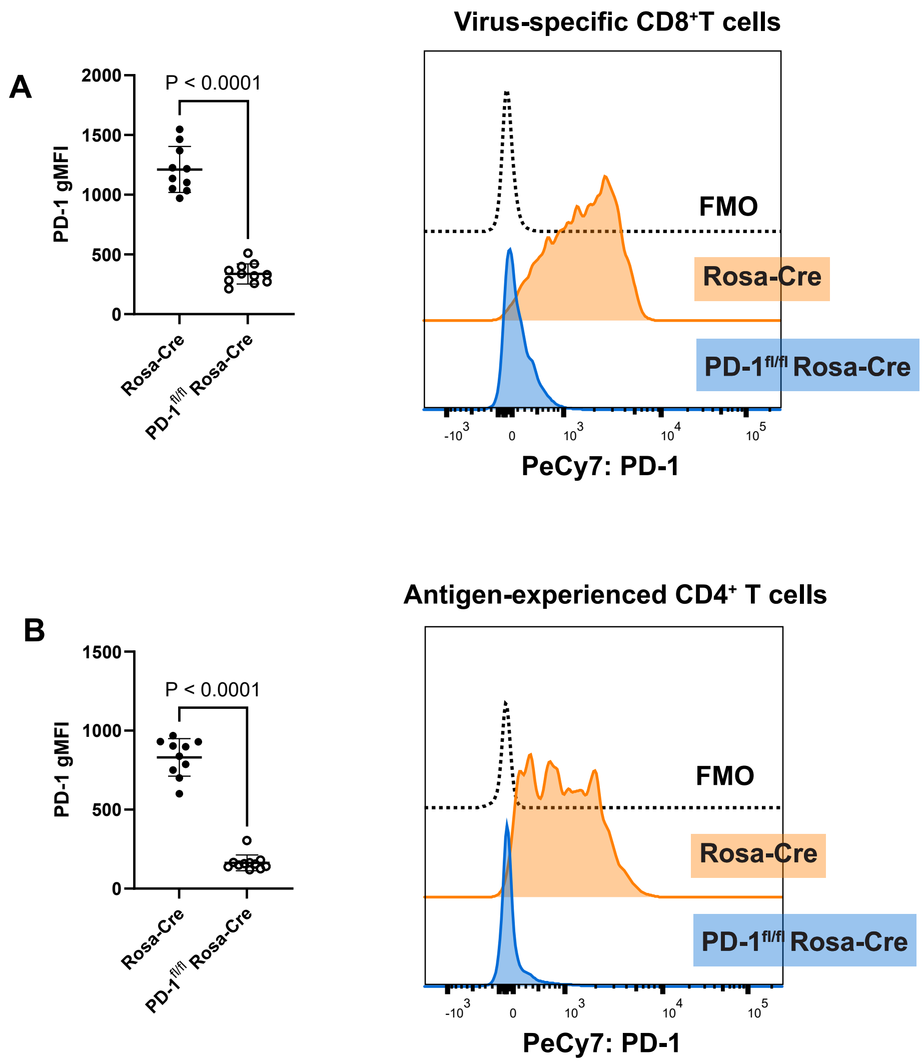


**Supplementary 3. Tamoxifen-induced knockdown of PD-1.** Rosa-Cre controls and PD-$1^{fl/fl}$ Rosa-Cre mice were infected with MuPyV i.c., given tamoxifen 8 to 12 dpi, and brains dissected at 15 dpi. **A)** PD-1 expression on brain-infiltrating virus-specific $CD8^+$ T cells. **B)** PD-1 expression on brain-infiltrating antigen-experienced $CD4^+$ T cells. n = 10 Rosa-Cre mice and 11 PD-$1^{fl/fl}$ Rosa-Cre mice. Error bars are mean ± SD. Data show two combined independent experiments. Statistical significance was determined with a Mann-Whitney U test.

### Supplementary 4

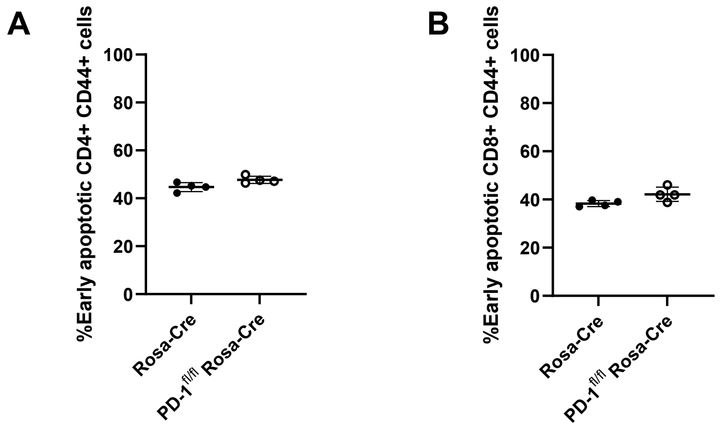


**Supplementary 4. Loss of PD-1 does not impact T cell apoptosis.** Rosa-Cre mice and PD-1$^{fl/fl}$ Rosa-Cre mice were treated as in Supplementary 3. Cells were assessed for early apoptosis via positive staining for annexin V and negative expression of viability dye, which stains dead cells. **A)** Frequency of antigen-experienced CD4$^{+}$ T cells undergoing early apoptosis. **B)** Frequency of virus-specific CD8$^{+}$ T cells going through early apoptosis. n = 4 Rosa-Cre mice and 4 PD-1$^{fl/fl}$ Rosa-Cre mice. Statistical significance was calculated with a Mann-Whitney U test. Error bars indicate mean ± SD.

### Supplementary 5

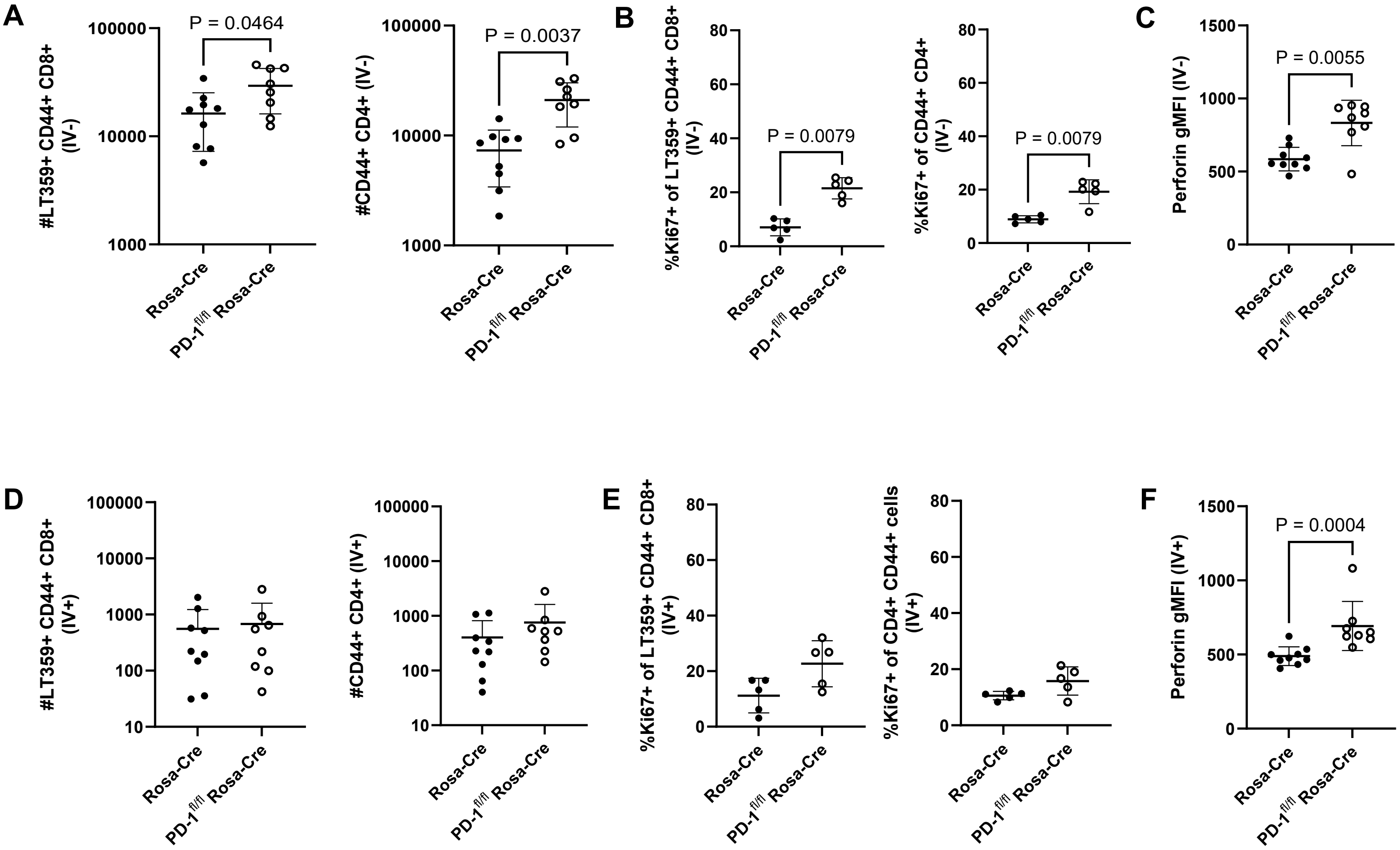


**Supplementary 5. Brain T cell phenotyping and quantification after CD45 antibody IV labeling.** Rosa-Cre mice and PD-1$^{fl/fl}$ Rosa-Cre mice were treated as in Supplementary 3 and immediately prior to euthanasia were injected IV with fluorophore-conjugated anti-CD45. **A)** Numbers of brain-infiltrating (IV-) virus-specific CD8$^+$ and antigen-experienced CD4$^+$ T cells. **B)** Proportions of IV- T cells positive for proliferation marker Ki67. **C)** Perforin expression in IV- virus-specific CD8$^+$ T cells. **D)** Numbers of T cells in the intravascular (IV+) compartment. **E)** Frequency of T cells expressing Ki67 in the IV+ compartment. **F)** Perforin expression in IV+ virus-specific CD8+ T cells. n = 9 Rosa-Cre mice and 8 PD-1$^{fl/fl}$ Rosa-Cre mice. Error bars are mean ± SD. Data combined from two independent experiments for all except B) and E), which are representative of two independent experiments. Statistical significance for all graphs was calculated with a Mann-Whitney U test.

### Supplementary 6

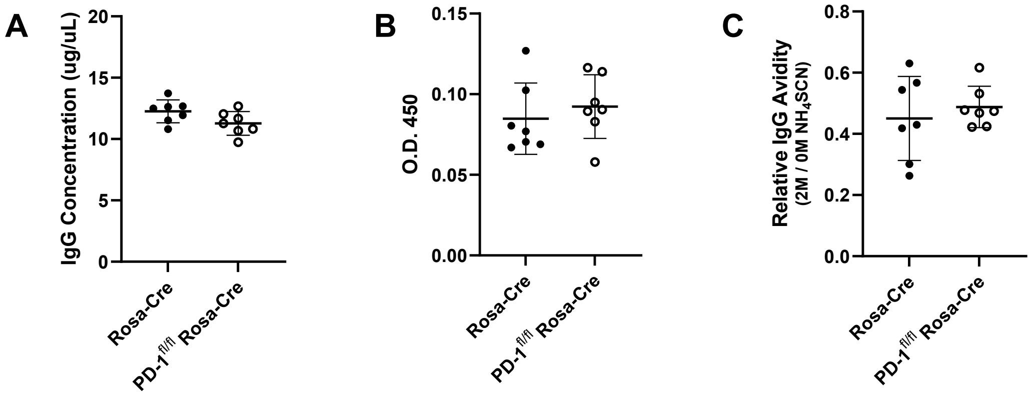


**Supplementary 6. PD-1 does not affect antiviral IgG or IgM.** Sera were collected from Rosa-Cre and PD-1$^{fl/fl}$ Rosa-Cre treated as in Supplementary 3, and IgG and IgM were quantified by ELISA. **A)** Concentration of anti-MuPyV IgG. **B)** IgM levels, shown by absorption. **C)** Relative IgG avidity to MuPyV. n = 7 Rosa-Cre mice and 7 PD-1$^{fl/fl}$ Rosa-Cre mice. Data combined from two independent experiments. Statistical significance quantified with a Mann-Whitney U test. Error bars show mean ± SD.

### Supplementary 7

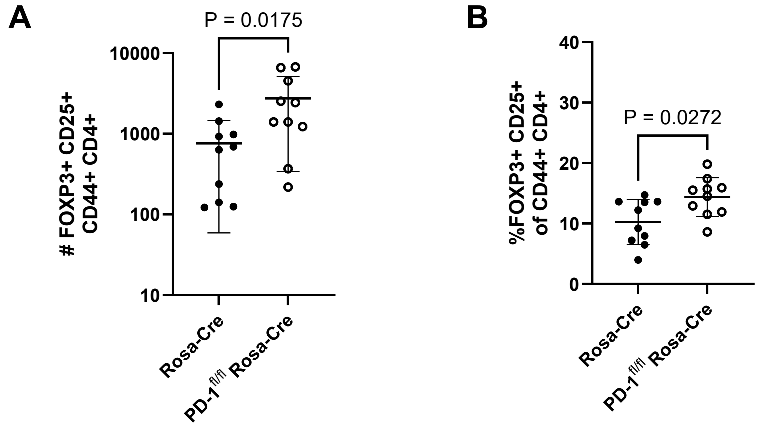


**Supplementary 7. PD-1 loss modestly increases $T_{reg}$ cells in brains of MuPyV-infected mice.** Mice were treated as in Supplementary 3. **A)** Numbers of $T_{reg}$ cells, designated as $FOXP3^+$ $CD25^+$ $CD44^+$ $CD4^+$. **B)** Proportion of antigen-experienced $CD4^+$ T cells positive for FOXP3 and CD25. n = 10 Rosa-Cre mice and 10 PD-$1^{fl/fl}$ Rosa-Cre mice. Error bars show mean ± SD. Data combined from two independent experiments. Statistical significance calculated with a Mann-Whitney U test.

### Supplementary 8

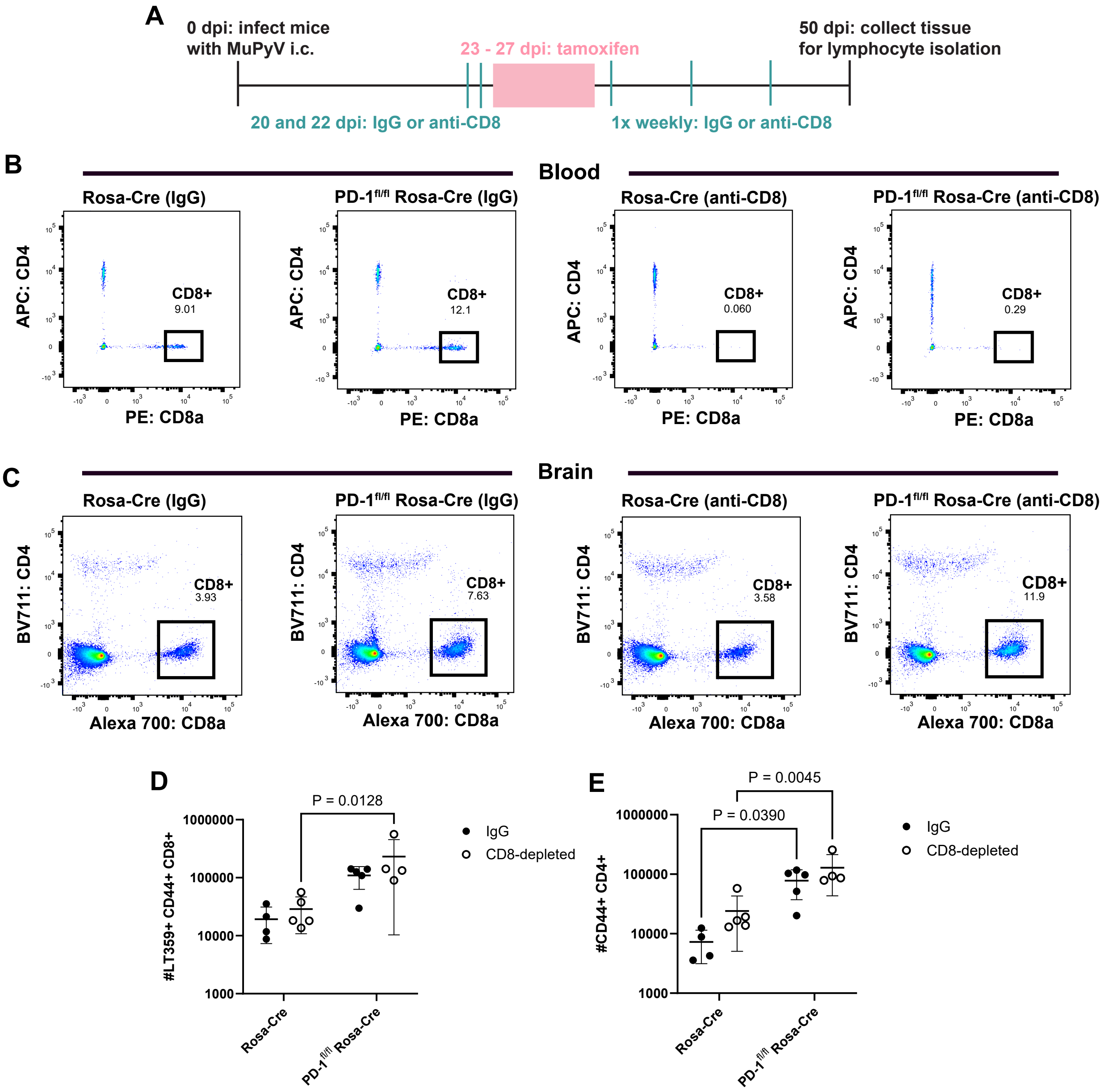


**Supplementary 8. T cells expand in a brain-autonomous manner after loss of PD-1 during persistent infection. A)** Experimental setup for antibody-mediated peripheral $CD8^+$ T cell depletion. Rosa-Cre and PD-$1^{fl/fl}$ Rosa-Cre mice were given either IgG control or anti-CD8 antibody prior to tamoxifen administration, then given either IgG or anti-CD8 once a week until endpoint. **B)** Representative flow cytometry plots showing $CD8^+$ and $CD4^+$ T cells in blood of IgG-treated and anti-CD8-treated mice. **C)** Representative flow cytometry plots showing $CD8^+$ and $CD4^+$ T cells in brains of IgG-treated and anti-CD8-treated mice. B) and C) were gated from live $CD45^+$ cells. **D)** Virus-specific $CD8^+$ T cell and **E)** $CD4^+$ T cell counts. n = 4 Rosa-Cre mice (rat IgG), 5 Rosa-Cre mice (anti-CD8), 5 PD-$1^{fl/fl}$ Rosa-Cre (rat IgG), and 4 PD-$1^{fl/fl}$ Rosa-Cre (anti-CD8). Statistical significance for D) and E) was determined with a two-way ANOVA with uncorrected Fisher's LSD post-test. Error bars indicate mean ± SD.

### Supplementary 9

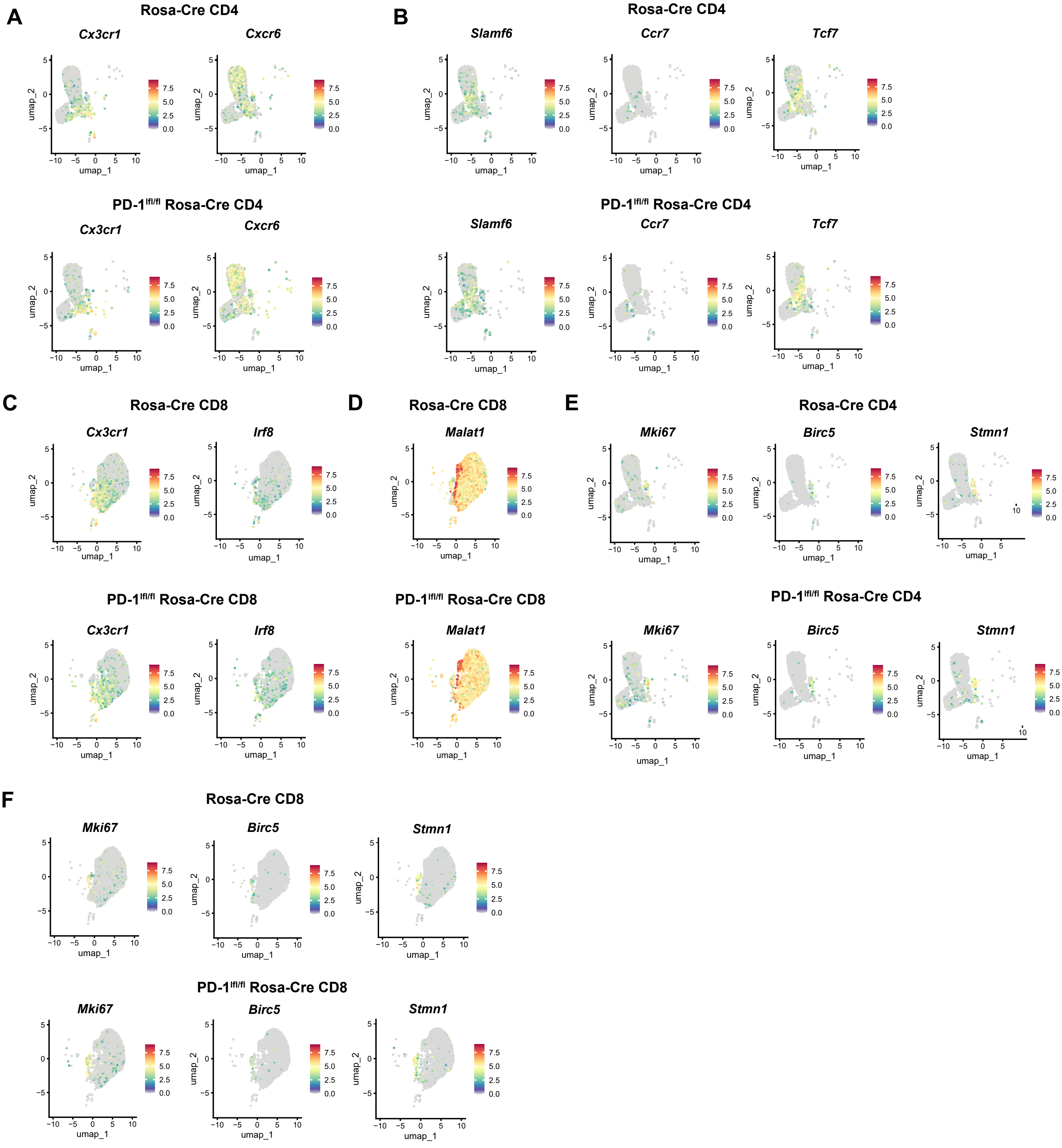


**Supplementary 9. Feature plots of genes in CD4⁺ and CD8⁺ T cell clusters.** Virus-specific CD8⁺ T cells and antigen-experienced CD4⁺ T cells were sorted from pooled groups of Rosa-Cre and PD-1$^{fl/fl}$ Rosa-Cre mice, and processed for scRNA-seq. **A)** Feature plots for *Cx3cr1* and *Cxcr6* in antigen-experienced CD4⁺ T cells. **B)** Feature plots for genes in cluster 6 of CD4⁺ T cells. **C)** Genes in cluster 0 of virus-specific CD8⁺ T cells. **D)** Feature plots for *Malat1* in cluster 8 of CD8⁺ T cells. **E)** Feature plots of genes in cluster 7 in CD4⁺ T cells. **F)** Feature plots of genes in cluster 7 in CD8⁺ T cells. n = 10 mice per strain.

### Supplementary 10

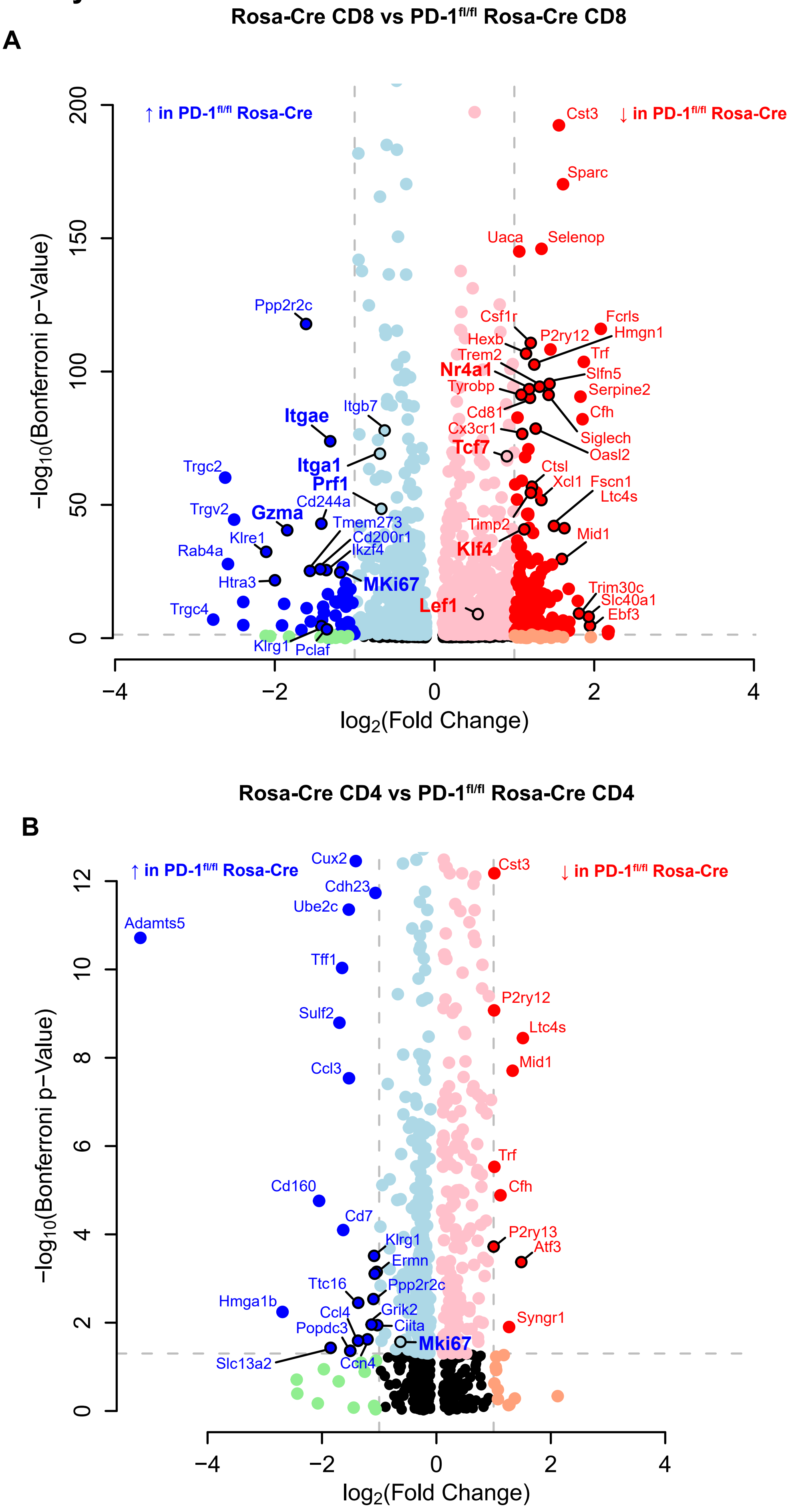


**Supplementary 10. Volcano plots showing differentially expressed genes in CD8⁺ and CD4⁺ T cells.** Genes in blue are upregulated with loss of PD-1, and genes in red are downregulated with loss of PD-1. Bolded genes indicate ones associated with effector, resident memory-like, or proliferative phenotypes. **A)** Differentially expressed genes in CD8⁺ T cells. **B)** Differentially expressed genes in CD4⁺ T cells. n = 10 per mice. Thresholds set at log2 fold change below -1 and above 1 and above -log10(0.05).

### Supplementary 11

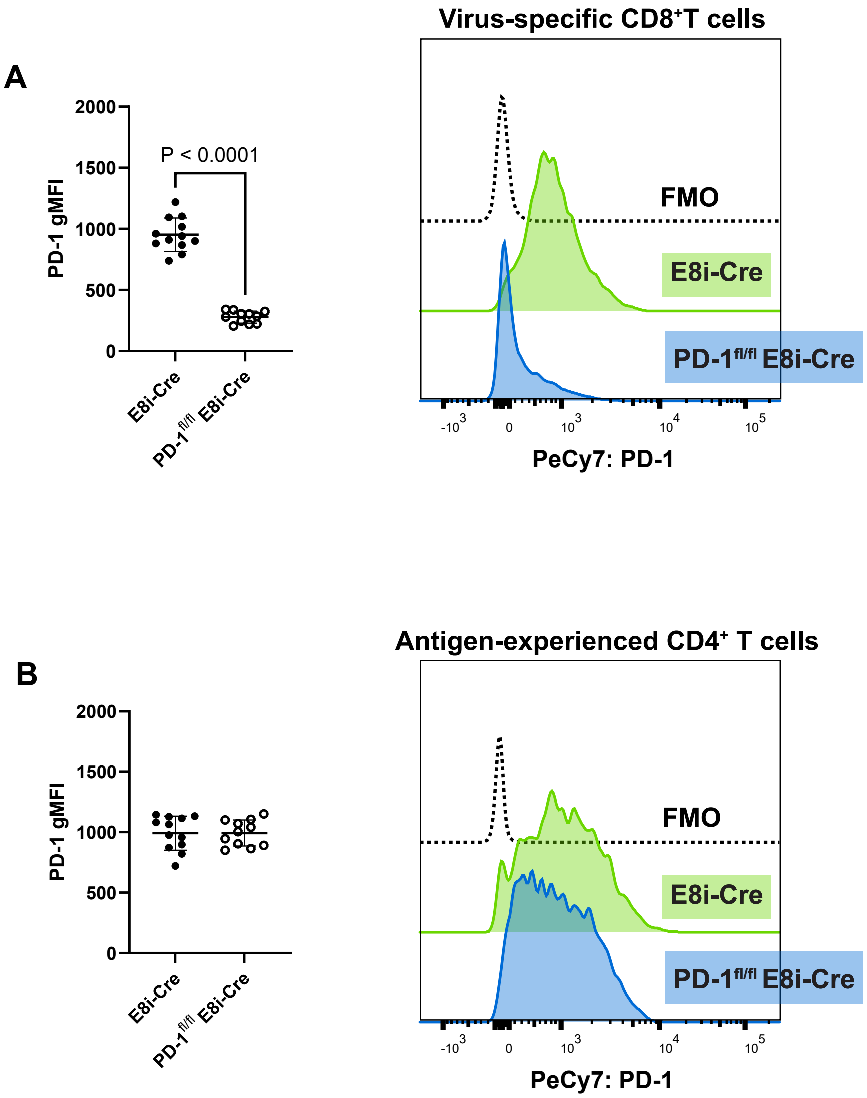


**Supplementary 11. CD8⁺ T cell-specific PD-1 knockdown.** E8i-Cre controls and PD-1$^{fl/fl}$ E8i-Cre mice were i.c. inoculated with MuPyV, administered tamoxifen at 8 to 12 dpi, and T cells isolated from brains at 15 dpi. **A)** PD-1 expression on virus-specific CD8⁺ T cells. **B)** PD-1 expression on antigen-experienced CD4⁺ T cells. n = 12 E8i-Cre mice and 11 PD-1$^{fl/fl}$ E8i-Cre mice. Data combined from two independent experiments. Statistical significance calculated with a Mann-Whitney U test.

### Supplementary 12

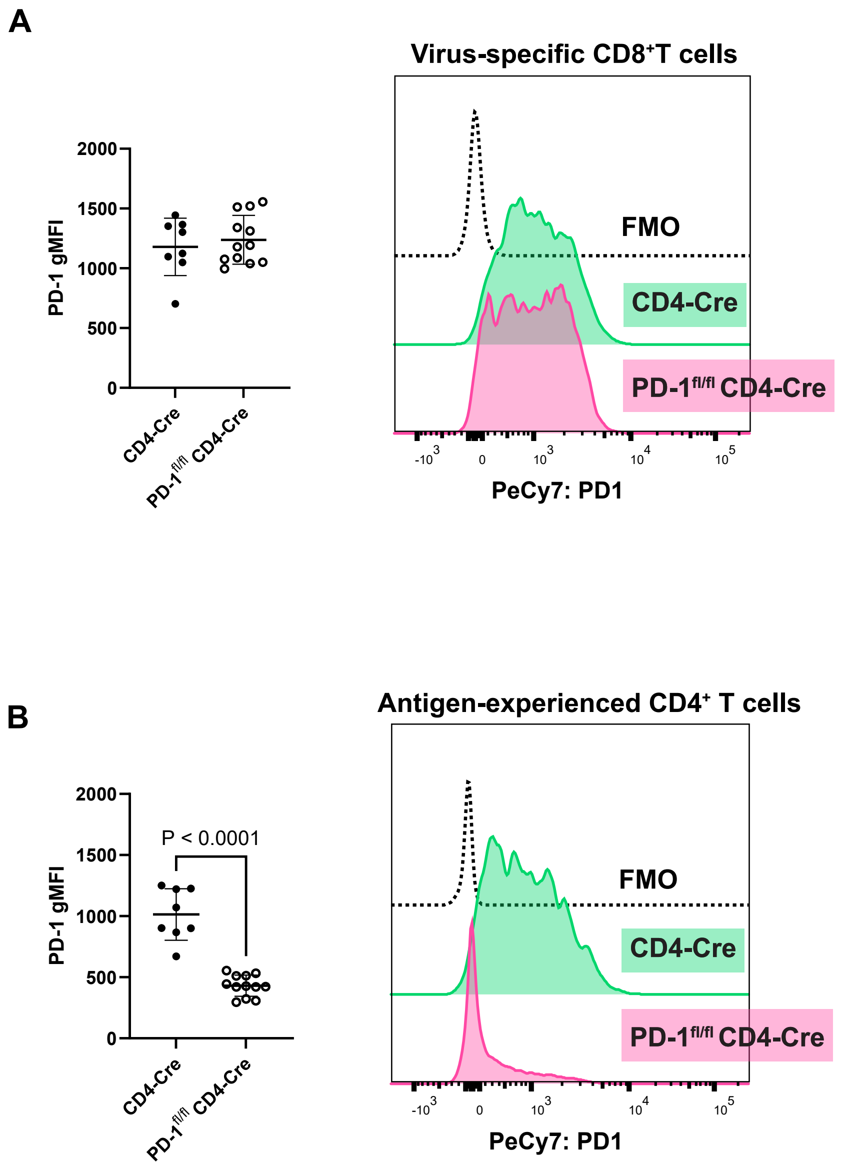


**Supplementary 12. CD4+ T cell-specific PD-1 knockdown.** CD4-Cre mice and PD-$1^{fl/fl}$ CD4-Cre mice were infected with MuPyV i.c. and given tamoxifen at 8 to 12 dpi. Brains were phenotyped through flow cytometry at 15 dpi. **A)** PD-1 expression on virus-specific $CD8^+$ T cells. **B)** PD-1 expression on antigen-experienced $CD4^+$ T cells. n = 8 CD4-Cre mice and 12 PD-$1^{fl/fl}$ CD4-Cre mice. Data combined from two independent experiments. Statistical significance quantified with a Mann-Whitney U test.
