## Supplemental Table 1 for "PD-1 regulates CD4^+^ T cell-mediated CD8^+^ T cell responses in the brain to balance viral control and neuroinflammation"

**Supplementary Table 1: List of Reagents or Resources used in Methods.**

| REAGENT or RESOURCE | SOURCE | IDENTIFIER |
|---|---|---|
| **Antibodies** | | |
| Anti CD8a PE (Clone 53.6-7) | Biolegend | Cat. 100708 |
| Anti-annexin V PE | Biolegend | Cat. 640908 |
| Anti-CD103 BV480 (Clone M290) | BD Biosciences | Cat. 566118 |
| Anti-Cd11b BV480 (Clone M1/70) | BD Biosciences | Cat. 566149 |
| Anti-CD19 FITC (clone 1D3/CD19) | Biolegend | Cat. 152404 |
| Anti-CD25 PerCPCy5.5 (Clone PC61) | Biolegend | Cat. 102029 |
| Anti-CD3 PE (Clone 500A2) | Biolegend | Cat. 152310 |
| Anti-CD4 APC (Clone RM4-5) | Biolegend | Cat. 100516 |
| Anti-CD4 BV650 (Clone RM4-5) | Biolegend | Cat. 100546 |
| Anti-CD4 BV711 (Clone RM4-5) | BD Biosciences | Cat. 563726 |
| Anti-CD44 BV785 (Clone IM7) | Biolegend | Cat. 103059 |
| Anti-CD45 AF700 (Clone 30.F11) | Biolegend | Cat. 103128 |
| Anti-CD45 BV605 (Clone 30-F11) | Biolegend | Cat. 103155 |
| Anti-CD45 FITC (Clone 30.F11) | Biolegend | Cat. 103108 |
| Anti-CD45 PerCPCy5.5 (Clone 30.F11) | Biolegend | Cat. 103132 |
| Anti-CD8a AF700 (Clone 53.6-7) | Biolegend | Cat. 100730 |
| Anti-CD8b (Clone H35-17.2) | Golstein et al., 1982 | N/A |
| Anti-FOXP3 AF700 (Clone MF-14) | Biolegend | Cat. 126422 |
| Anti-Granzyme B Pacific Blue$^{TM}$ (Clone GB11) | Biolegend | Cat. 515408 |
| Anti-IFNy (APC) | BD Biosciences | Cat. 554413 |
| Anti-IgM biotin | Biolegend | Cat. 406504 |
| Anti-MHCII (I-A/I-E) APC (Clone M5/114.15.2) | Biolegend | Cat. 107614 |
| Anti-NK1.1 BV605 (Clone PK136) | Biolegend | Cat. 108739 |
| Anti-PD-1 PeCy7 (Clone RMP1-30) | Biolegend | Cat. 109110 |
| Anti-PDL1 BV421 (Clone MIH5) | BD Biosciences | Cat. 564716 |
| Anti-Perforin PE (Clone S16009A) | Biolegend | Cat. 154306 |
| Anti-TCF1 FITC (Clone 63D9) | Cell Signaling | Cat. 6444S |
| ChromPure Rat IgG | Jackson Immunoresearch | Cat#012-000-003 |
| eBioscience™ Anti-Ki67 AF700 (Clone SolA15) | Thermo Fisher Scientific Invitrogen$^{TM}$ | Ref. 56-5698-82 |
| Goat anti-Mouse IgG Heavy and Light Chain Antibody HRP Conjugated | Bethyl Laboratories | Ref: A90-116P |
| HRP Goat Anti-IgG (minimal x-reactivity) Antibody | Biolegend | Ref: 405405 |

| | | |
|---|---|---|
| HRP Streptavidin | Biolegend | Cat. 405210 |
| LT359 APC | NIH Tetramer | Core/RRID:SCR_026557 |
| **Virus strains** | | |
| MuPyV, Strain A2 | N/A | N/A |
| **Chemicals, peptides, and recombinant proteins** | | |
| BD GolgiPlug™ Protein Transport Inhibitor (Containing Brefeldin A) | BD Biosciences | Cat. 51-2301KZ |
| Collagenase (Type I) | Worthington | Cat. LS004197 |
| Corn oil | Sigma-Aldrich | C8267 |
| DNAse I | Worthington | Cat. LS002140 |
| Percoll | Cytiva | Product 17089101 |
| Tamoxifen | Sigma-Aldrich | T5648 |
| **Commercial assays** | | |
| 1-Step™ TMB ELISA Substrate Solutions | Thermo Fisher Scientific | Ref. 34029 |
| BD Rhapsody WTA Reagent Kit - 8 Pack | BD Biosciences | Cat. 666620 |
| BD Rhapsody™ Cartridge Kit | BD Biosciences | Cat. 633733 |
| Bioanalyzer High Sensitivity DNA Analysis RUO | Agilent | Part No. 5067-4626 |
| eBioscience™ Foxp3 / Transcription Factor Staining Buffer Set | Thermo Fisher Scientific Invitrogen | Cat. 00-5523-00 |
| EIA/RIA Polystyrene High Bind Microplate | Corning | Ref. 3590 |
| Ms Single Cell Sample Multiplexing Kit | BD Biosciences | Cat. 633793 |
| PerfeCTa FastMix II ROX | QuantaBio | Part No. 95119-012 |
| Wizard Genomic DNA Purification Kit | Promega | Ref. A1120 |
| **Experimental model organisms** | | |
| C57Bl/6J | Jackson Laboratories | Strain No. 000664 |
| CD4-CreERT2 C57BL/6J | Jackson Laboratories | Strain No. 022356 |
| E8i-Cre C57BL/6J | Andrews et al., 2021 | N/A |
| PD-1fl/fl C57BL/6J | Laurent Brossay at Brown University | N/A |
| Rosa26 CreERT2 | Jackson Laboratories | Strain No. 004847 |
| **Software** | | |
| BCL Convert | Illumina | N/A |
| Benchling | Benchling | RRID:SCR_013955 |
| Flowjo | BD Biosciences | RRID:SCR_008520 |
| Graphpad Prism | Dotmatics | RRID:SCR_002798 |
| R v. 4.4.3 | R Core Team 2021 | RRID:SCR_001905 |
| **Other** | | |

| | | |
|---|---|---|
| BD Pharmingen™ Annexin V Binding Buffer, 10X concentrate | BD Biosciences | Cat. 556454 |
| BD Omics-Guard Sample Preservation Buffer | BD Biosciences | Cat. 570911 |
| eBioscience™ Fixable Viability Dye eFluor™ 780 | Thermo Fisher Scientific Invitrogen™ | Cat. 65-0865-14 |
